# Preventing post-surgical cardiac adhesions with a catechol-functionalized oxime hydrogel

**DOI:** 10.1101/2020.12.29.424755

**Authors:** Masaki Fujita, Gina M. Policastro, Austin Burdick, Hillary T. Lam, Jessica Ungerleider, Rebecca L. Braden, Diane Huang, Kent Osborn, Jeffery H. Omens, Michael M. Madani, Karen L. Christman

## Abstract

Post-surgical cardiac adhesions represent a significant problem during routine cardiothoracic procedures. This fibrous tissue can impair heart function and inhibit surgical access in reoperation procedures. Here, we propose a novel hydrogel barrier composed of oxime crosslinked poly(ethylene glycol) (PEG) with the inclusion of a catechol (Cat) group to improve retention on the heart for pericardial adhesion prevention. This three component system is comprised of aldehyde (Ald), aminooxy (AO), and Cat functionalized PEG mixed to form the final gel (Ald-AO-Cat). Ald-AO-Cat has favorable mechanical properties, degradation kinetics, and minimal swelling, as well as superior tissue retention compared to an initial Ald-AO gel formulation. We show that the material is cytocompatible, resists cell adhesion, and led to a reduction in the severity of adhesion in an *in vivo* rat model and a pilot porcine study. The Ald-AO-Cat hydrogel barrier may therefore serve as a promising solution for preventing post-surgical cardiac adhesions.

---

Depressed fibrinolytic activity resulting from surgical trauma to the epicardium during open heart procedures, can lead to fibrous adhesion formations between the epicardium and other tissues in the chest cavity.^1–3^ Formation of these fibrous adhesions impedes heart function and severely complicates resternotomy by obstructing visibility and increasing the risk of mortality and morbidity during dissection.^2,4–6^ For children born with congenital heart defects, who will experience multiple surgeries over their lifetime, and adults receiving valve replacements, mechanical circulatory support, and/or coronary artery bypass grafting, this problem is particularly relevant.^2,7,8^ Reoperations for adult patients now constitute more than 20% of the annual caseload in cardiac surgery.^9^

A variety of materials have been studied for the reduction and prevention of surgical adhesions, such as Seprafilm (sodium hyaluronate/carboxymethylcellulose sheet) and CoSeal (polyethylene glycol (PEG) hydrogel).^10–15^ However, limited success has been demonstrated for preventing or reducing the severity of cardiac adhesions due to short retention time as a result of post-surgical swelling and dynamic motion of the heart, short degradation times, and excessive swelling of the polymer leading to cardiac tamponade.^10-12,14-18^ One product was approved in the U.S. for preventing cardiac adhesions, REPEL-CV (polylactic acid/PEG sheet), although this failed to reduce adhesion dissection time^19^ and is no longer sold.

A promising method for ameliorating cardiac adhesion formation is to coat the epicardium with a fast-gelling polymer barrier to prevent susceptible tissue from adhering to the sternum or other organs in the chest cavity.^4,12,17^ *In situ* polymerization/crosslinking of hydrogels after spraying has been tested for many adhesion prevention applications using a variety of different chemistries, including thiol-Michael addition reactions, “click” chemistry, amine-aldehyde imine-forming reactions, and photoinitiated radical crosslinking.^20–26^ Designing these polymer coatings for the heart is, however, challenging, because the polymer material must: 1) be easily applied; 2) rapidly gel on the tissue surface in an aqueous environment; 3) be retained on the epicardium for at least 2 weeks to overcome the initial inflammatory response post-surgery; 4) exhibit biocompatibility; and 5) have minimal swelling to prevent cardiac tamponade.^4,13,14,27^ Ideally, the polymer could also be easily removed if a surgeon requires immediate re-access to the heart in case of emergency.

Our earlier studies identified an oxime-crosslinked star PEG-based injectable hydrogel consisting of aldehyde terminated PEG and aminooxy terminated PEG (Ald-AO) as a new approach to potentially prevent post-operative pericardial adhesions.^28^ PEG-based hydrogel systems are resistant to protein and cell adhesion, and oxime bonds form rapidly and have excellent hydrolytic stability.^28–31^ Unlike other hydrogel systems with high swelling ratios (~400 %) that have been studied for cardiac anti-adhesion applications, this Ald-AO PEG-oxime hydrogel rapidly forms gels with high crosslink density, allowing for a reduced swelling ratio, which should minimize the risk of cardiac tamponade when translated *in vivo*.^14,32^ Despite the ability of aldehydes to bind to amines on the epicardial surface, preliminary *in vivo* studies in a rat model showed that retention of the material on the heart was not optimal, which led to inconsistent results with adhesion prevention (unpublished). Herein, we have developed a novel three-component PEG-based, injectable oxime hydrogel system (Ald-AO-Cat), utilizing the mussel-inspired catechol (Cat) found on dopamine (DA) for prolonged retention time on cardiac tissue and prevention of adhesion formation (Figure 1A). Catechol compounds, such as DA, participate in strong covalent and non-covalent interactions with both organic and inorganic substrates. Particularly, mussel adhesive proteins are enriched in 3,4-dihydroxy-phenylalanine (DOPA), which is attributed to its ability to bind strongly to surfaces under water.^33–35^ When exposed to air, catecholic compounds are oxidized to quinones which further react with nucleophiles, such as amines.^36–38^ Catechol chemistry has been studied for many biomedical applications, particularly as tissue adhesive materials, with much success,^39–44^ which motivated our use of it to improve polymer retention on the heart. Interestingly, in one study using a catechol-functionalized PEG hydrogel for islet transplantation in a murine model, researchers made note of the complete absence of non-specific adhesions to the implant, which was attributed to the high PEG content of the hydrogel.^45^ However, to the best of our knowledge, a catechol-conjugated hydrogel has never been utilized in a material designed to be applied to an organ to prevent post-surgical adhesions.

**Figure 1.**
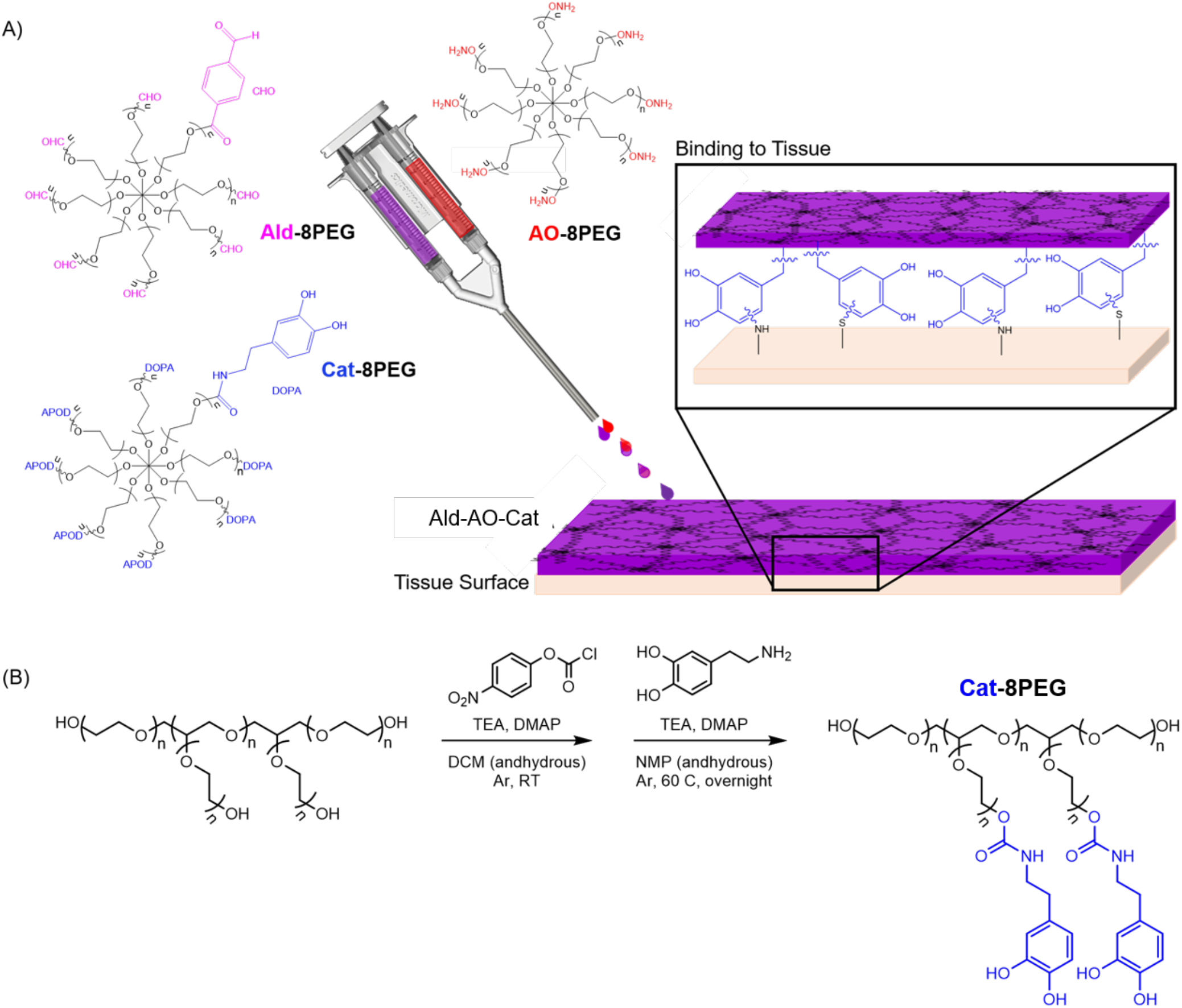
A) Three distinct 8-arm star PEG polymers were used to form the new oxime hydrogel. The Ald-8PEG/Cat-8PEG solution is mixed with the AO-8PEG solution in a 1:1 ratio and sprayed onto the tissue surface using a FibriJet gas-assisted applicator head from Nordson Micromedics. Oxime bonds rapidly form between Ald-8PEG and AO-8PEG. The catechol forms covalent bonds with the primary amines present on the tissue surface to promote hydrogel-tissue adhesion. (B) Synthesis of Cat-8PEG required a two-step process by activating PEG-OH with NPC and then deprotecting with dopamine hydrochloride. Ald-8PEG and AO-8PEG were synthesized as previously described.^46^ Abbreviations: triethylamine (TEA), 4-(dimethylamino) pyridine (DMAP), dichloromethane (DCM), N-methyl-2-pyrrolidone (NMP), 4-nitrophenyl chloroformate (NPC).

Herein, we tested the use of the Ald-AO-Cat gel for specifically preventing cardiac post-surgical adhesions through studying its mechanical and biological properties via parallel-plate rheometric, retention assays, and cellular adhesion over time. Additionally, this study demonstrates the cytocompatibility of the hydrogel through metabolic assays performed after direct contact with an *in vitro* cell layer. Lastly, the Ald-AO-Cat system was implanted into small and large animal cardiac adhesions models to investigate its performance on the surface of the heart. We have demonstrated the use of this hydrogel barrier system in reducing the severity and preventing cardiac adhesions in a rat model of post-surgical cardiac adhesions as well as in a pilot porcine study, indicating its promising clinical application in cardiothoracic procedures.

## Results

### Synthesis and characterization of polymers

This novel oxime-hydrogel system required the synthesis of three distinct 8-arm PEG polymers: aldehyde-functionalized (Ald-8PEG), aminooxy-functionalized (AO-8PEG) and catechol-functionalized (Cat-8PEG). Ald-8PEG and AO-8PEG were synthesized as previously described.^47^ Cat-8PEG was synthesized in two steps by activation of 8-arm PEG-OH with 4-nitrophenyl chloroformate (NPC) followed by deprotection with dopamine hydrochloride (Figure 1B). ^1^H NMR (Figure S1-S3) was used to confirm the successful synthesis and the functionalization ratio of each PEG component. Catechol-functionalization was further verified using UV-Vis (Figure S4) because of the hydroscopic nature of PEG and the overlap of H_2_O and PEG proton peaks in the ^1^H NMR. The average number of functionalized arms were calculated and are reported in Table 1. Functionalization ratios of polymers ranged from 74-97 %, which can be equated to the successful functionalization of 5.9-7.8 arms of the PEG-OH polymer for each PEG polymer.

**Table 1.** Polymer functionalization.

| Polymer | Functionalization<sup>a</sup><br>(%) | Functionalization<sup>b</sup><br>(%) | Number of Arms<br>Functionalized |
| --- | --- | --- | --- |
| Ald-8PEG | 97 | --- | 7.8 |
| AO-8PEG | 83 | --- | 6.6 |
| Cat-8PEG | 74 | 81 | 5.9<sup>a</sup> (6.5)<sup>b</sup> |

<sup>a</sup> Calculated from <sup>1</sup>H NMR peak integrations
<sup>b</sup> Calculated from UV-Vis

### Formation and Characterization of Hydrogels

All hydrogels were formed with a fixed 1:1 ratio of Ald-8PEG: AO-8PEG because excess aldehyde groups significantly affected the metabolic activity of 3T3 fibroblasts in a larger elution extract as previously reported.^47^ All gel formulations are listed in Table 2.

**Table 2.**
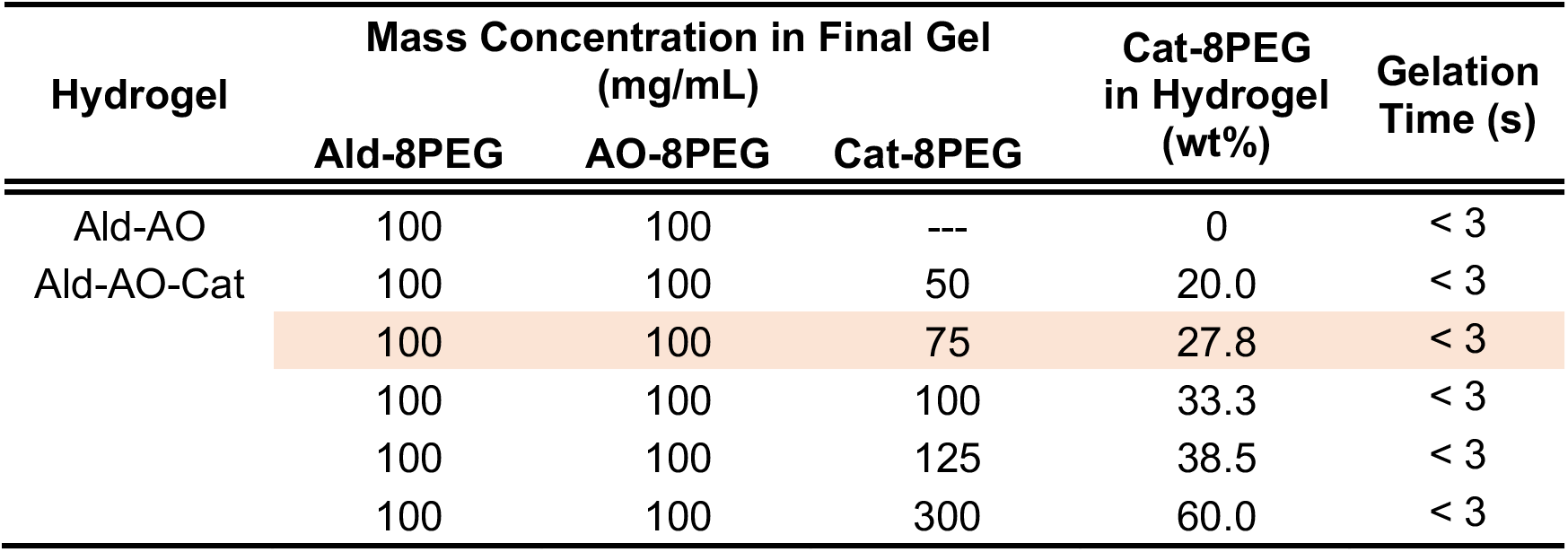
Hydrogel formulations.

Storage modulus (G’) for Ald-AO gels was studied as a function of polymer concentration (Figure 2A). G’ values ranged from 3.0 ± 0.3 kPa to 20.1 ± 1.2 kPa for 25 mg/mL and 150 mg/mL of total polymer, respectively. For 100 mg/mL, G’ was 17.0 ± 0.6 kPa; this hydrogel polymer concentration was chosen for the Ald-AO gel as well as for further testing with incorporation of Cat-8PEG. The ratio of Cat-8PEG was varied for Ald-AO-Cat gel formation to find the optimal hydrogel parameters. All formulations gelled rapidly (< 3 s) regardless of Cat-8PEG content (Table 2). Increased Cat-8PEG content in Ald-AO-Cat gels, resulted in a decreased G’ (Figure 2B), ranging from 17.0 ± 0.6 kPa to 2.6 ± 1.8 kPa for 0 wt% Cat-8PEG and 50 wt% Cat-8PEG incorporation, respectively. The catechol-oxime hydrogel consisting of 100 mg/mL of Ald-8PEG and AO-8PEG and 75 mg/mL (27.8%) of Cat-8PEG (G’ = 8.1 ± 1.0 kPa) was evaluated in subsequent studies. Relative G’ was calculated at 1 and 3 hours, compared to G’ at the initial gel formation (Figure 2C). G’ significantly increases from 0 to 3 hours for Ald-AO-Cat, compared to Ald-AO, which ultimately shows no change in G’. At 3 hours post gelation, G’ for Ald-AO-Cat increased 150 ± 20 %. A representative frequency sweep of G’ and G” exhibits the viscoelastic behavior of the crosslinked gels (Figure S5). Rheometry was also used to calculate the individual polymer solution viscosities, which were 4.2 x 10^-2^ Pa·s and 1.0 × 10^−2^ Pa·s for Ald/Cat-8PEG and AO-8PEG solutions, respectively (Figure S7).

**Figure 2.**
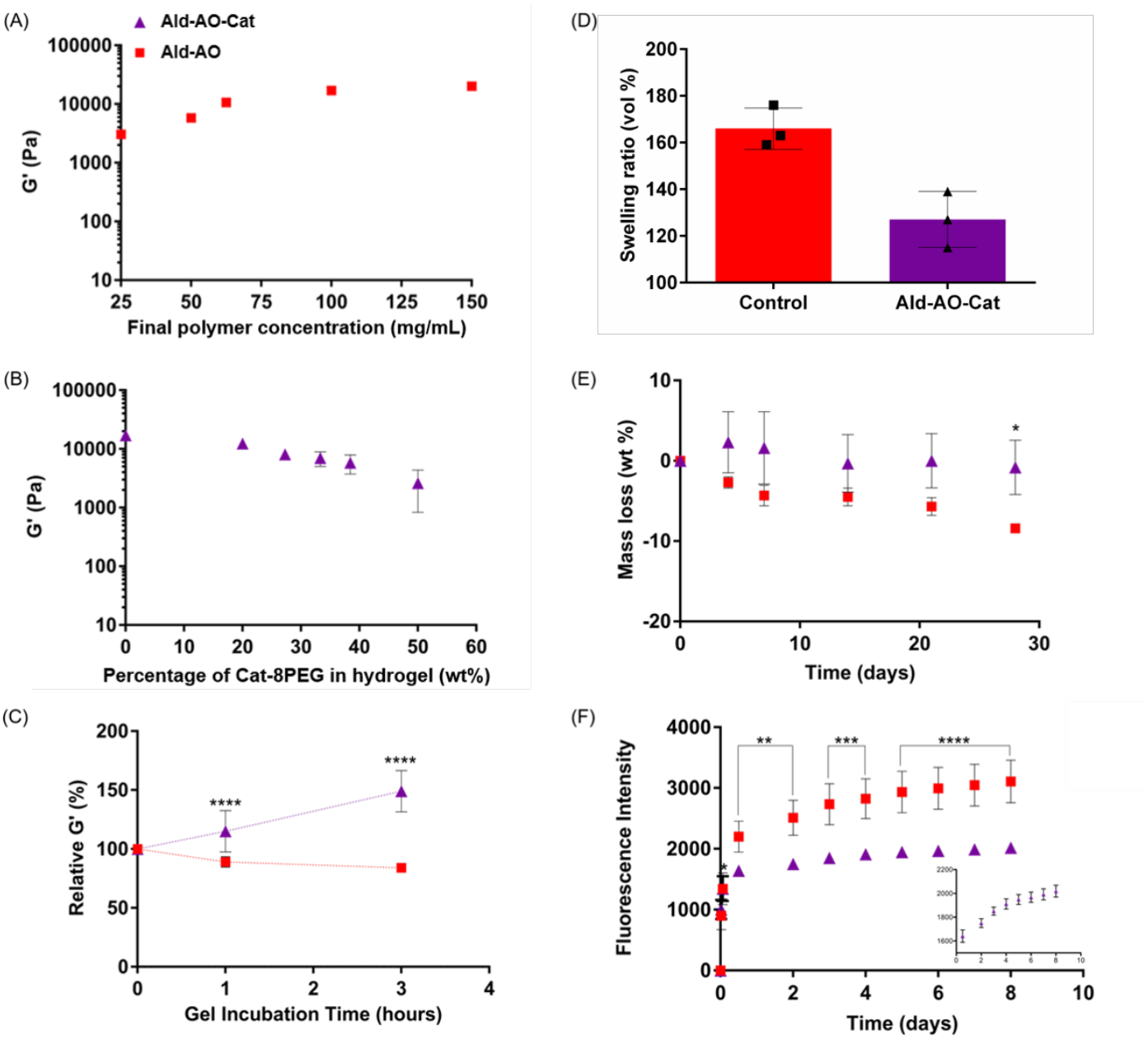
Characterization of rapidly-gelling oxime hydrogels. (A) Storage moduli (G’) were measured as a function of final polymer mass concentration in Ald-AO gels using a frequency sweep from 10^-2^ to 10^2^ Hz at 37 °C; G’ at 1 Hz is shown. (B) Ald-AO-Cat gels with varying Cat-8PEG content were gelled and G’ was measured under the same conditions. Increasing Cat-8PEG content of the gels resulted in decreasing G’. (C) G’ for Ald-AO-Cat and Ald-AO were measured up to 3 hours post gelation and relative G’ was calculated. Over the 3 hour incubation period, G’ significantly increased for Ald-AO-Cat indicating the formation of secondary crosslinks between Cat and AO. (D) Low swelling ratios were observed for both Ald-AO-Cat and Ald-AO gels, with significantly lower swelling ratio observed for Ald-AO-Cat (E) In vitro degradation of the oxime hydrogels revealed significantly greater mass loss after 28 days for Ald-AO gels compared to Ald-AO-Cat gels. (F) *Ex vivo* retention of Ald-AO-Cat shows significantly greater retention on the tissue surface compared to Ald-AO over 8 days. Inset shows Ald-AO-Cat only to allow visualization of the error bars. Mean ± SD and analyzed with an unpaired t-test (swelling ratio) or a one-way ANOVA with a Tukey’s post-hoc test (*p < 0.05, **p < 0.005, ***p < 0.001, ****p < 0.0001). n=3 for each condition.

The Ald-AO-Cat and Ald-AO gels exhibited 127 ± 12% and 166 ± 9% swelling, respectively, in PBS (pH 7.4) after reaching swelling equilibrium in 24 h (Figure 2D). Degradation of Ald-AO-Cat and Ald-AO gels was tested *in vitro* for 28 days in PBS (pH 7.4). A mass loss of 8.4 ± 0.4 wt% was observed for Ald-AO gel, which was significantly greater than the 0.8 ± 3.4 wt% mass loss observed for Ald-AO-Cat after 28 days (Figure 2E).

Retention of the oxime hydrogel systems was tested in PBS (pH 7.4) on *ex vivo* porcine cardiac biopsy punches using fluorescently labeled Ald-8PEG and AO-8PEG (Figure 2F). For both the Ald-AO and Ald-AO-Cat gels, a burst release of polymer was observed within the first 12 h. However, after 2 days, significantly less fluorescently labeled polymer was released from Ald-AO-Cat gels, compared to the Ald-AO gel. This pattern was observed through 8 days. Additionally, the retention of Cat-8PEG was observed in a time-dependent release study over 7 days. Similar to the *ex vivo* retention data, a burst release of Cat-8PEG was seen in the day 1 supernatant, with decreasing concentration of Cat-8PEG released overtime (Figure S6A). The total amount of Cat-8PEG released and the remaining Cat-8PEG in the gel after 7 days of incubation was compared to the total Cat-8PEG content in the gel formulation at the time of gelation. About 37 % of the Cat-8PEG was not accounted for using this method, which is attributed to the incomplete digestion of the Cat-8PEG gel at day 7, resulting in additional Cat-8PEG sticking in the gel pellet and not being detected with the assay reading (Figure S6B).

### *In vitro* cell studies

L929 fibroblast and RAW macrophage adhesion to Ald-AO-Cat and Ald-AO gels was compared to control tissue culture (TC) plastic adhesion (Figure 3A). Cells were seeded on each surface and fluorescent images of prelabeled cells were taken to determine percent surface coverage after 24 h incubation. Less than 1% fluorescent area was observed for both cell lines on hydrogel surfaces, while 6.5 ± 1.5% and 22.0± 3.3% fluorescent area was observed for RAW macrophages and L929 fibroblasts, respectively, on TC plastic controls.

**Figure 3.**
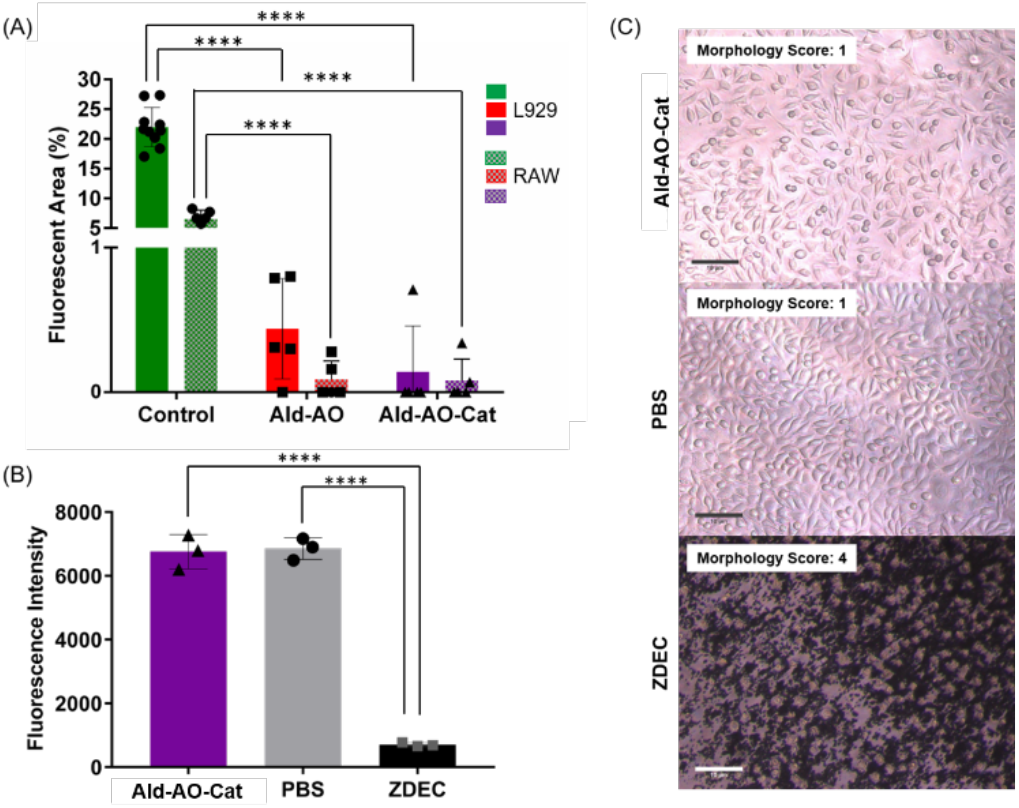
In vitro testing of Ald-AO-Cat. (A) Inflammatory cell adhesion was studied on oxime hydrogels and tissue culture plastic as a control. L929 fibroblasts and RAW macrophages were seeded on each surface and imaged after 24 h of incubation for cell adhesion, as indicated by fluorescent area. Less than 1% fluorescent area was observed for both gels for L929 and RAW cells. Conversely, 22% and 6.6% fluorescence area were observed for L929 and RAW, indicating oxime hydrogels resist inflammatory cell adhesion. (B) Cytocompatibility of Ald-AO-Cat gels was assessed after 24 h of incubation in direct contact with the L929 fibroblast cell monolayer. PBS and ZDEC were doped into the media for positive and negative controls, respectively. Ald-AO-Cat did not significantly affect metabolic activity compared to the positive control. (C) Brightfield images of the fibroblasts in each condition reveal that no effect on cell morphology and spreading was observed for Ald-AO-Cat compared to the positive control. A cell morphology score was assigned to each brightfield image for quantification of cell morphology. Morphology scores suggest that fibroblasts are well spread and that no apparent cell lysis occurs when in direct contact with Ald-AO-Cat. Data are reported as Mean ± SD and analyzed with a one-way ANOVA with a Tukey’s post-hoc test (****p < 0.0001). n=3 for each condition.

Cytocompatibility of the Ald-AO-Cat gels was tested with a direct contact assay by forming the Ald-AO-Cat gel directly on an L929 fibroblast monolayer. Cells were incubated for 24 h and cytocompatibility was assessed via metabolic activity (Figure 3B) and imaging (Figure 3C) against a positive and negative control. After 24 h incubation, alamar blue readings indicate that there was no effect on metabolic activity in the presence of Ald-AO-Cat. Resulting fluorescent intensities showed no significant difference from PBS controls (6760 ± 538 compared to 6855 ± 340, respectively). Brightfield images were obtained for assessing cell morphology for each treatment group (Figure 3C). Images were blinded from the grader and morphology scores ranging from 1-4 were assigned. Morphology scores are defined in Table 3 in the methods section. When treated with Ald-AO-Cat and PBS, cells consistently received a score of 1 (≤ 20% of cells appear rounded with minimal lysing), while ZDEC control wells were given scores of 4 (All cells appear rounded and lysed). Both the alamar blue and morphology scores indicate the cytocompatibility of Ald-AO-Cat.

**Table 3.**
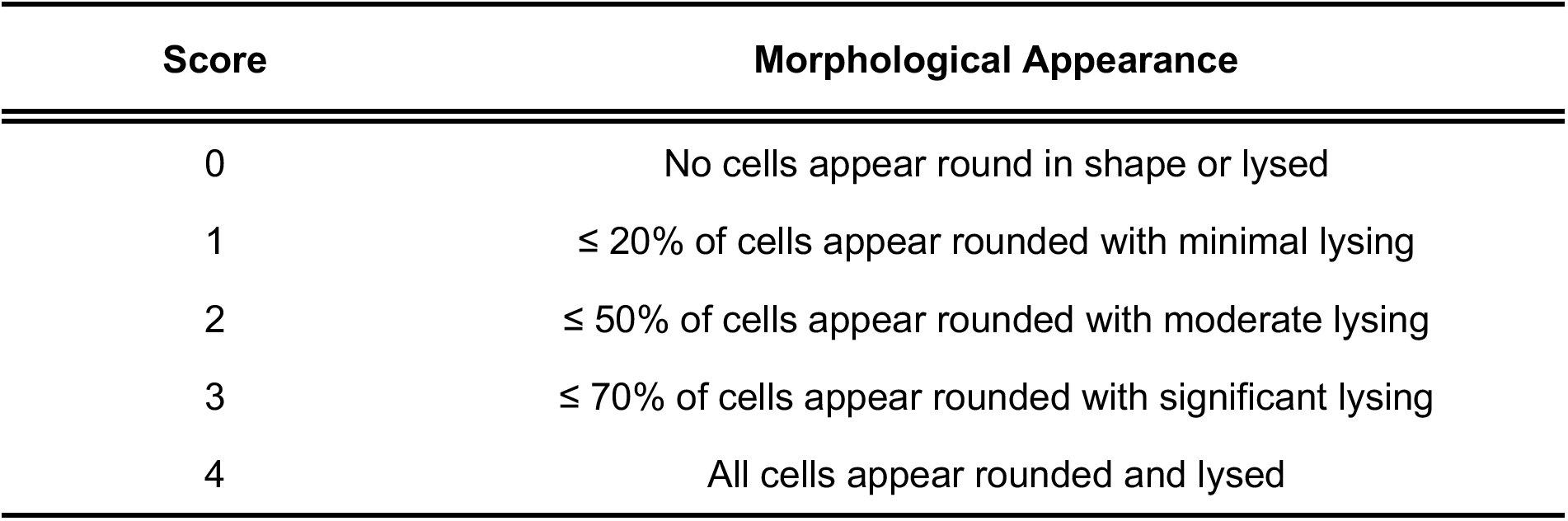
*in vitro* cell morphology scores.

| Score | Morphological Appearance |
| --- | --- |
| 0 | No cells appear round in shape or lysed |
| 1 | ≤ 20% of cells appear rounded with minimal lysing |
| 2 | ≤ 50% of cells appear rounded with moderate lysing |
| 3 | ≤ 70% of cells appear rounded with significant lysing |
| 4 | All cells appear rounded and lysed |

### *In vivo* rat cardiac adhesion model

A rat cardiac adhesion model was used to test the ability of the Ald-AO-Cat gel to prevent cardiac adhesions compared to Ald-AO gel and untreated rats. Gross assessment of cardiac adhesion formation was performed when the chest was re-entered. Before dissecting the adhesions to harvest the heart, images were taken and given to blinded graders for assigning adhesion scores. The heart was divided into 9 segments, and each segment was given an overall adhesion score based on the presence and severity of adhesions from 0-4 (Figure 4A). The average score over these 9 segments was reported as the average adhesion score. The average adhesion intensity was calculated over all regions that showed adhesion formation. The maximum adhesion intensity score was also reported for each animal. This is a common method used for adhesion scoring in various *in vivo* models.^48–50^ Representative images of the observed adhesions in control and Ald-AO-Cat groups are shown in Figure 4A.

**Figure 4.**
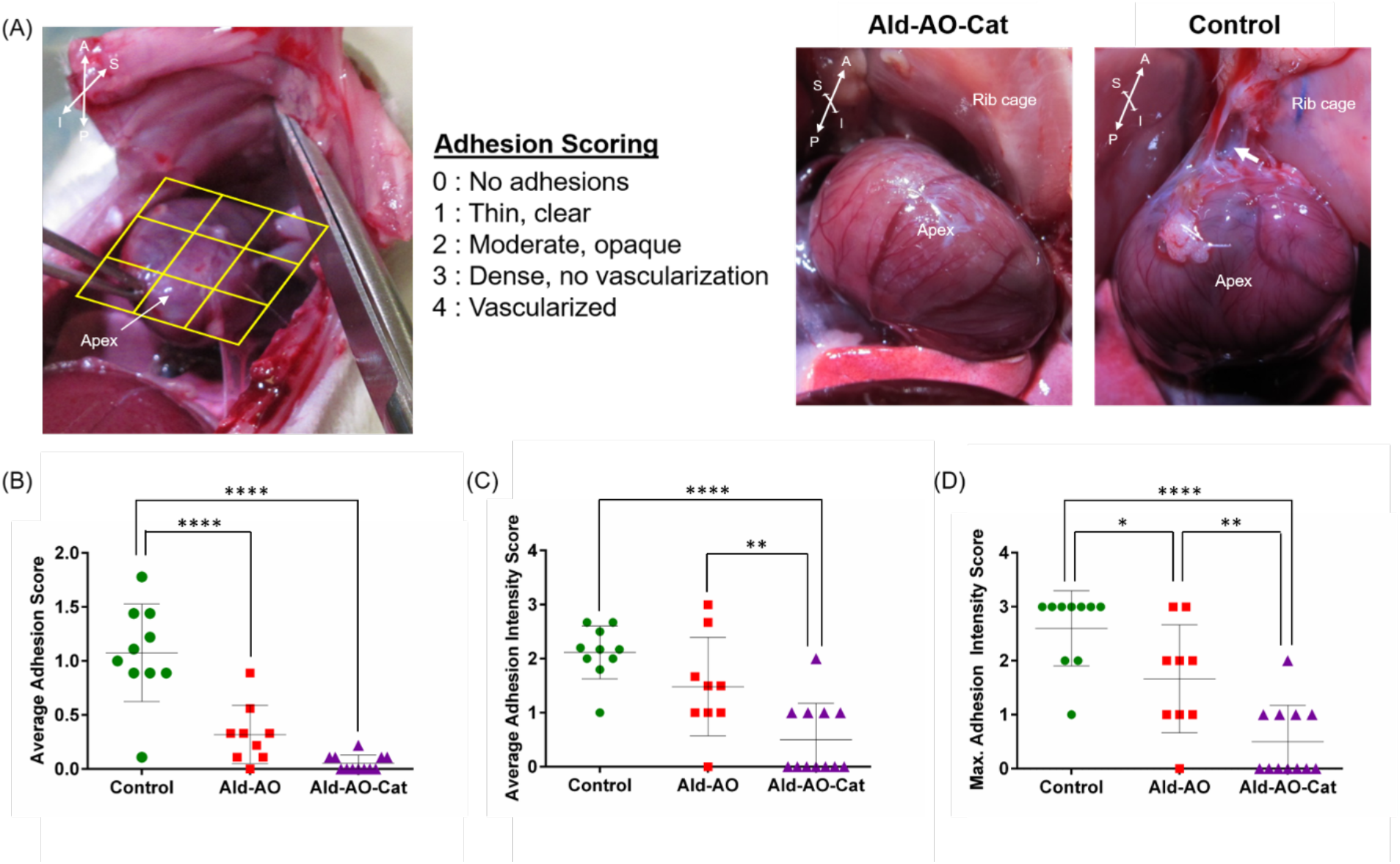
Adhesion formation and heart function were assessed for administration of Ald-AO-Cat, Ald-AO and the untreated control after 2 weeks. (A) Adhesion scores were assigned to 9 discrete segments of the exposed heart surface when the chest was re-entered during euthanasia and harvest. Representative images of adhesions upon re-entry for Ald-AO-Cat and control groups are shown. Anatomical directions are shown to provide a point of reference: A-anterior, P-posterior, S-superior, I-inferior (B) The individual scores for the 9 segments were averaged to obtain the adhesion score for each heart. At 2 weeks there was a significant reduction in adhesion score for Ald-AO-Cat and Ald-AO compared to the untreated group. (C) The average intensity score was calculated over regions of adhesion formation. At 2 weeks, Ald-AO-Cat show significantly reduced adhesion intensity compared to Ald-AO and the untreated control. (D) The maximum adhesion intensity score was also reported for each animal. Significantly higher maximum intensity scores were assigned to adhesions in rats treated with Ald-AO and the untreated group, compared to Ald-AO-Cat. Together these data suggest reduced adhesion formation and intensity when Ald-AO-Cat is applied following abrasion. Data are reported as Mean ± SD and analyzed with a one-way ANOVA with a Tukey’s post-hoc test (*p < 0.05) n = 12, 9, and 10 for Ald-AO-Cat, Ald-AO and Control groups, respectively.

At 2 weeks, there was a significant reduction in average adhesion score when Ald-AO-Cat and Ald-AO were applied, compared to the untreated control (Figure 4B). All of the treatment groups consistently received low adhesion scores, however Ald-AO-Cat (0.1 ± 0.1) and Ald-AO (0.3 ± 0.3) groups were significantly lower compared to the untreated (1.1 ± 0.5) control at 2 weeks. The average intensity scores of the adhesions showed similar results, with a significantly reduced adhesion intensity score when Ald-AO-Cat (0.5 ± 0.7) was applied, compare to Ald-AO (1.5 ± 0.9) and the untreated control (2.1 ± 0.5). There was no difference in average adhesion intensity score reported for Ald-AO and untreated groups. Maximum adhesion intensity score was the final parameter used to grade adhesion formation and hydrogel efficacy. Compared to Ald-AO (1.7 ± 1.0) and untreated groups (2.6 ± 0.7), Ald-AO-Cat (0.5 ± 0.7) application resulted in significantly lower maximum adhesion intensity.

Adhesion formation and cardiac function were assessed in a second *in vivo* study, comparing untreated rats to Ald-AO-Cat, since it was shown to be superior to the Ald-AO gel in the first study. At 4 weeks, a reduction in adhesion formation and severity was observed (Figure 5). All of the rats treated with Ald-AO-Cat showed no signs of adhesion formation and received an overall adhesion score of 0, which was significantly lower than the untreated group (0.2 ± 0.2) (Figure 5A). Consistent with 2 week results, the average adhesion intensity score (Figure 5B) and maximum adhesion intensity score (Figure 5C) for Ald-AO-Cat (scores of 0 for both measurements) were also significantly lower than the untreated group (average adhesion intensity 0.9 ± 0.8), maximum adhesion intensity 1.0 ± 0.9).

**Figure 5.**
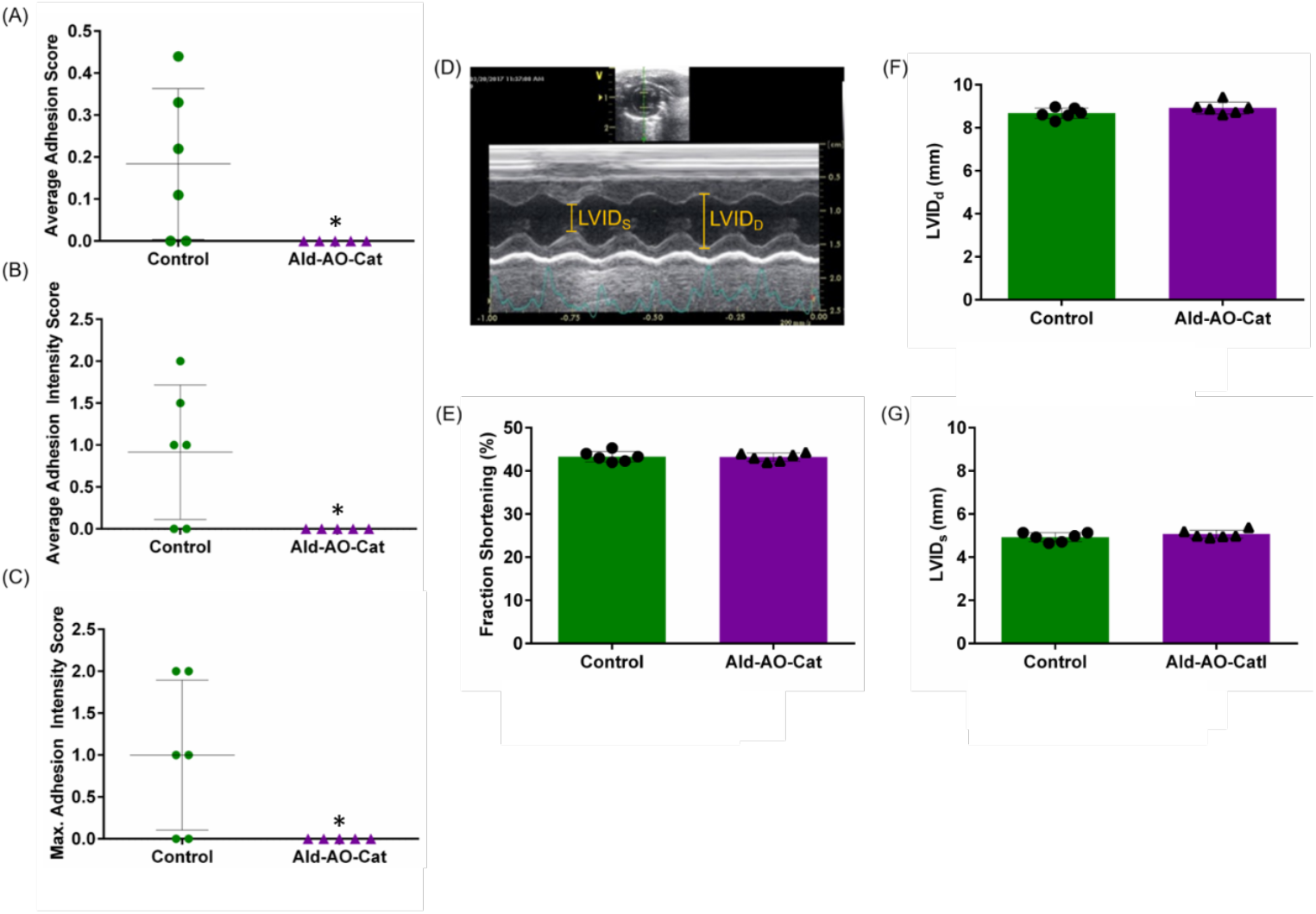
Adhesion formation and heart function were assessed for administration of an untreated control and Ald-AO-Cat. (A-C) At 4 weeks, all parameters for comparing adhesion formation and intensity showed significantly lower values for Ald-AO-Cat when compared to the untreated control. It is also worth noting that all Ald-AO-Cat treated animals received an average adhesion and intensity score of 0, indicating complete reduction of adhesion formation. (D-G) Cardiac function was assessed 3 ± 1 days post material application using M-mode echocardiography. Fraction shortening was calculated, and left ventricle internal diameter systole (LVID_S_) and left ventricle internal diameter diastole (LVID_D_) were measured, and regardless of treatment, animals showed identical cardiac functioning, indicating no adverse effects from application of Ald-AO-Cat gels. Data are reported as Mean ± SD and analyzed with an unpaired Mann-Whitney t-test (*p < 0.05). n = 5 and 6 for Ald-AO-Cat and Control groups, respectively.

To ensure the hydrogels did not impede cardiac function, M-mode echocardiography was conducted 3 ± 1 days post material application (Figure 5D-G). Three days was chosen to allow for full swelling *in vivo* to ensure that cardiac tamponade would not occur. When Ald-AO-Cat was applied there was no difference in end-diastolic left ventricular internal diameter (LVID_D_), end-systolic left ventricular internal diameter (LVID_S_), or fractional shortening (FS) compared to the untreated group, indicating normal cardiac function following gel application.

Histological assessment was performed by a trained histopathologist on hearts that were harvested at 4 weeks. Hematoxylin and Eosin (H&E) staining revealed the presence of some regions with Ald-AO-Cat hydrogel on the epicardium at 4 weeks, indicating that the material was not fully degraded. Minimal macrophage infiltration was visible at high magnification. In areas with patches of remaining hydrogel, there was evidence of some thin encapsulation (Figure 6), however there was no indication of lymphocyte and neutrophil infiltration, suggesting that no chronic inflammation was occurring and there was a resolved wound healing response.

**Figure 6.**
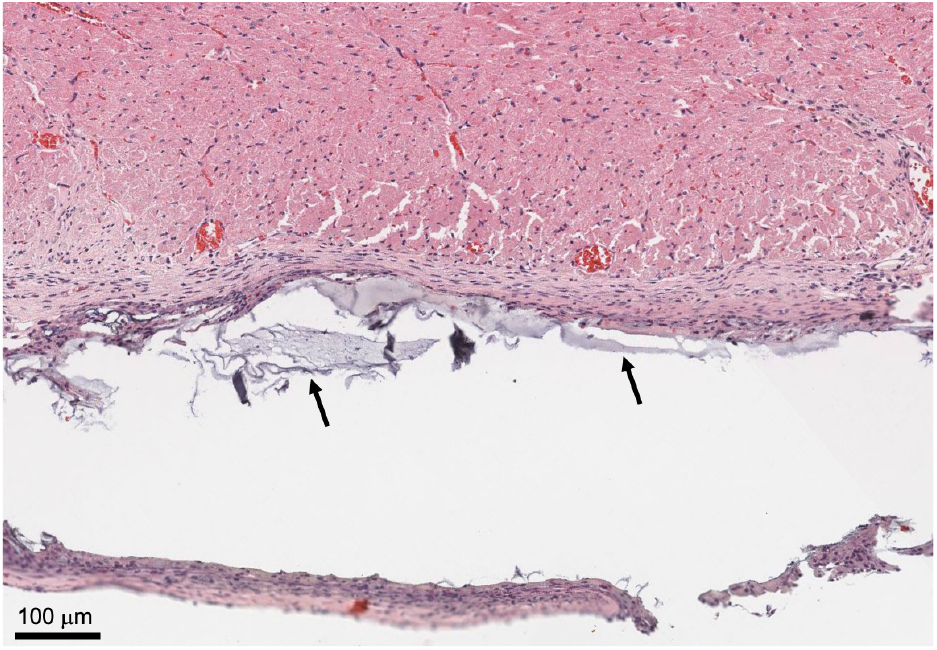
H&E staining was performed for Ald-AO-Cat tissue samples to verify biocompatibility of the oxime hydrogel systems. At 4 weeks, thin capsule formation around some remaining gel (black arrows) was visible, however there was no indication of chronic toxicity as a result of application or degradation of the hydrogel system. Furthermore, no cell infiltration into the Ald-AO-Cat gel coincides with our expectations, due to the noncell adhesive nature of PEG polymers.

### Dual-delivery device design and testing

For delivery of our hydrogel system in large animal model, an air-assisted dual delivery device was designed (Figure 7A). The device design allowed for optimal 1:1 delivery ratio of both Ald-Cat-8PEG and AO-8PEG solutions, over the entire anterior surface of the heart as confirmed by a colormetric spray test (Figure 7B). At all distances from the sprayed surface, the mixing ratio showed no significant difference compared to the standard 1:1 mixed solution. Spraying of the device with yellow and blue dyed PEG solutions onto epicardial tissue *ex vivo* resulted in a 0.5-1.0 mm thick homogenous gel on the tissue surface, which could be manually removed if necessary, with minimal overspray onto “surrounding tissue” when spraying from a working distance of 12-17 cm with 10 psi air pressure (Figure 7C-E).

**Figure 7.**
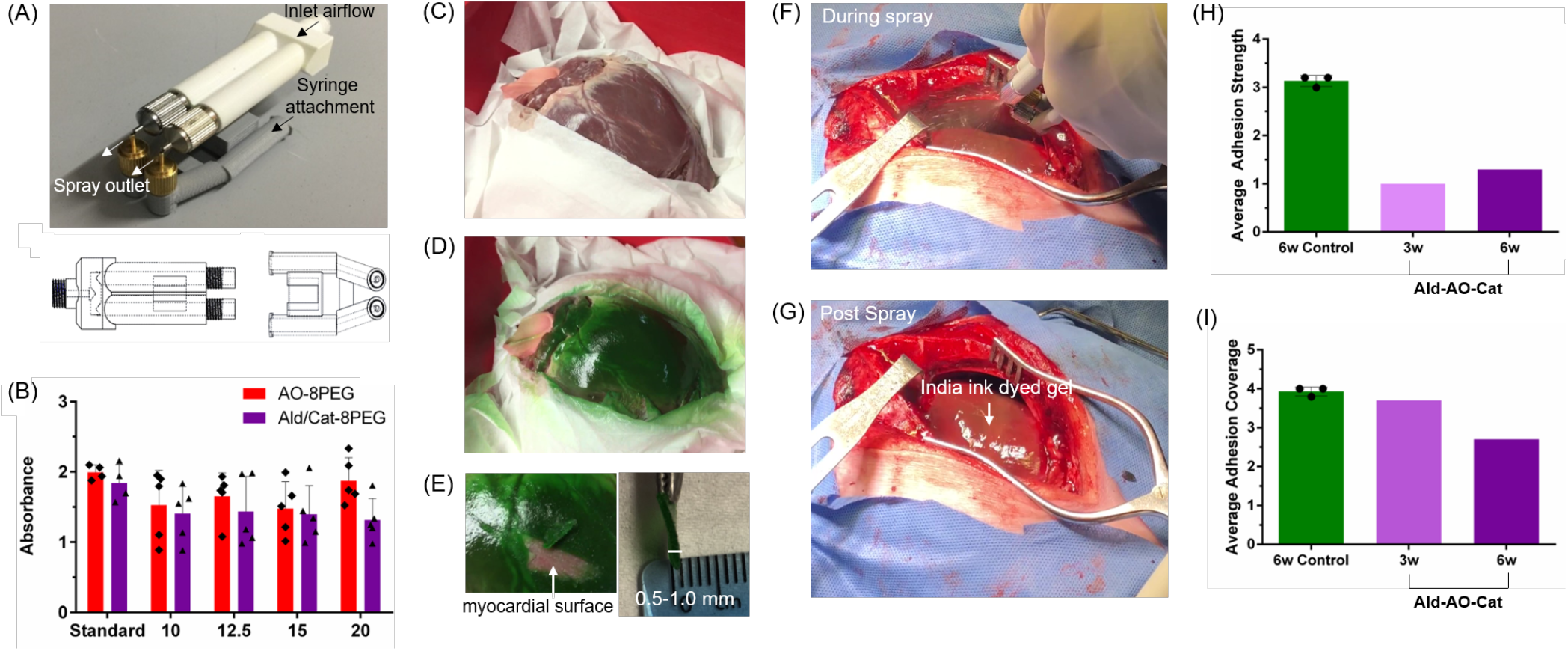
Spray device design, testing, and application in a pilot porcine study. (A) A specialized device was developed to deliver each component together in a 1:1 ratio in the form of an atomized spray for application in a large animal model. This device has two separate parts, one of which allows the flow of air from an air compressor through two separate pathways, while a second piece supports the addition of two syringes containing sterile solutions of the component polymers injected into the path of air through two pathways. (B) The mixing ratio of the sprayed solutions at various working distances was tested using mock Ald-Cat-8PEG and AO-8PEG solutions dyed blue and yellow, respectively. The standard was prepared by pipetting equal volumes of both solutions as a control (n=5). Equal mixing of both solutions was observed at all spray distances. (C) The dual spray device was tested on an *ex vivo* porcine heart to determine an effective working distance and the amount of spray necessary for complete coverage of the anterior surface with a gel ~0.5-1.0 mm thick. The heart was set in a shallow insulated ice bucket surrounded by kim wipes to simulate the exposure of the epicardium during a sternotomy. (D) The anterior surface of the heart was sprayed from a working distance of ~10-12 cm with a total of 8 mL (4 mL AO-8PEG (dyed blue) and 4 mL Ald-Cat-8EPG (dyed yellow)) of hydrogel solution. Equal mixing was visually determined by the green gel that formed on the surface of the heart. Minimal overspray is seen on the kim wipes surrounding the heart surface, indicating effective targeting of the spray. (E) For clinical translation, in the case of an emergency, it would be advantageous that the gel could be manually removed by the surgeon. The gel thickness was measured after removal, showing the targeted 0.5-1.0 mm gel thickness has been obtained with the 8 mL of solution. (F-G) The Ald-AO-Cat gel applied using the dual spray device was tested in a pilot porcine study against a non-treated control. The solutions were prepared with a 1:100 dilution of India Ink for visualization of the gel once sprayed. (H-I) 3 and 6 week post-sternotomy and abrasion, resternotomy and adhesion scoring was performed based on the defined adhesion scoring system (Table 4). Assessment of adhesions revealed a decrease in adhesion strength in Ald-AO-Cat treated animals (n=2), compared to controls (n=3). The treated animals required only simple blunt dissection of the adhesions to access the anterior surface of the heart, while the control required sharp dissection, resulting in tissue damage with adhesion removal. Average adhesion coverage was similar for all animals with a trending decrease in adhesion coverage at 6 weeks for the Ald-AO-Cat treated animal. This observation agrees with clinical observations of reduced cardiac adhesion coverage at longer time points post sternotomy and surgical trauma.

### *In vivo* porcine cardiac adhesion model pilot study

A porcine cardiac adhesions model was used to test the ability of Ald-AO-Cat to reduce the coverage and severity of cardiac adhesions compared to non-treated controls in a pilot investigation. Following a full sternotomy, the heart was abraded with gauze, and either treated with Ald-AO-Cat or left un-treated (Figure 7F-G). The heart surface was left exposed to air and the chest cavity was closed. Resternotomy and adhesion assessment was performed 3 weeks (n=1) and 6 weeks (n=1) post-gel application, and 6 weeks post-abrasion for all control animals (n=3) based on the scoring system defined in Table 4. The control pigs received an average adhesion coverage score of 3.9, while 3 weeks and 6 weeks Ald-AO-Cat treated pigs received scores of 3.7 and 2.6, respectively (Figure 7H). More importantly, despite the formation of adhesions in all animals, the control pig scored higher in adhesion strength (3.1) compared to 3 week and 6 week Ald-AO-Cat pigs (1.0 and 1.4, respectively) (Figure 7I). This semi-quantitative assessment of adhesion strength shows that most adhesions in the control pigs required sharp dissection and resulted in tissue damage when removed, while gel-treated pigs reduced the severity of adhesions which showed minimal fibrosis and required simple blunt dissection only. Representative images prior to dissection are shown in Figure S8.

**Table 4.** Pilot porcine adhesion scoring criteria.

| Score | Adhesion Coverage | Adhesion Strength |
| --- | --- | --- |
| 0 | 0 % | No adhesions |
| 1 | 1-25 % | Weak adhesions, remove with little force |
| 2 | 26-50 % | Moderately strong adhesions, blunt dissection required |
| 3 | 51-75 % | Strong adhesions, requires sharp dissection |
| 4 | 76-100% | Very strong adhesions, requires sharp dissection resulting in additional tissue damage |

## Discussion

In this study, oxime-crosslinked PEG hydrogels were synthesized and characterized for use as a cardiac anti-adhesion barrier. The oxime bond was chosen because previous work has demonstrated its rapid gel formation, chemo-specificity, aqueous stability and biocompatibility.^28,29,47,51^ The novel addition of Cat-8PEG was reported in this study, as the third component of this new oxime-hydrogel system for cardiac anti-adhesion applications. Inspired by nature, this polymer utilizes the adhesive properties of the catechol group in DA, a compound abundantly found in mussel proteins, which enables adhesion to a variety of surfaces in wet conditions.^52^ The hydroxyl groups of Cat-8PEG react with free amines,^38^ which exist on the epicardium, leading to an observed increased retention time on the tissue (Figure 2F).

All hydrogels were formed with a 1:1 ratio of Ald-8PEG:AO-8PEG, to ensure that unreacted aldehyde groups would not interfere with cell metabolic activity, as previously reported.^47^ While maintaining a fixed 1:1 ratio, the polymer concentration was varied to find the optimal formulation for Ald-AO gels. G’ is tunable with varying polymer concentration, with an observed increase in G’ with increasing polymer concentration. All polymer concentrations exhibited G’ within the range for cardiac muscle (~10-100 kPa), which is crucial for mimicking the elasticity of the native cardiac tissue.^53–56^ Gelation time (< 3 s) and the solution viscosities were taken into consideration when deciding the 100 mg/mL concentration would be the primary formulation for the Ald-AO gel. Tunable G’ was also observed when the concentration of Cat-8PEG was varied for the Ald-AO-Cat formulation. A trending decrease in G’ was observed with increase in Cat-8PEG wt%. The 27.8 wt% Ald-AO-Cat formulation was considered optimal because the solution viscosities did not complicate the delivery for gel formation, rapid gelation was achieved, and it resulted in a storage modulus that should not interfere with cardiac function. Overall, we wanted to use the highest wt% Cat-8PEG possible to ensure longer retention times on the tissue surface when applied *in vivo* without compromising ease of delivery through viscous solutions. Relative G’ measurements of Ald-AO-Cat resulted in an increase in G’ as incubation time increased, suggesting that physical entrapment of Cat-8PEG increases the storage modulus because of limited chain movement. This phenomenon was not observed for Ald-AO.

Swelling and degradation were used to characterize Ald-AO and Ald-AO-Cat gel formulations. At physiological pH in PBS there was minimal swelling (< 200 %) for both hydrogel systems. In clinical trials, the excess swelling (400 %) of CoSeal® was reported to be the cause of cardiac tamponade in several patients.^13,14^ Degradation of the both oxime hydrogel systems in PBS pH 7.4 over the course of 28 days revealed significantly greater degradation of the Ald-AO gel compared to Ald-AO-Cat. Slow degradation of Ald-AO-Cat revealed <1 % degradation over 28 days in physiological conditions.

Retention times of Ald-AO-Cat and Ald-AO were evaluated on *ex vivo* porcine biopsy punches over the course of 8 days. The goal was to develop a cardiac anti-adhesion barrier that would be retained on the native tissue for at least 2 weeks, to overcome the initial inflammatory response post-surgery.^57^ In preliminary unpublished work with the Ald-AO gel *in vivo,* we found this formulation had inadequate retention on the heart. In this study we sought to develop a material with better retention, which we hypothesized would form a more robust anti-adhesion barrier since several barrier materials have failed as cardiac anti-adhesion materials because of short retention and short degradation times.^10–12,15–17^ We demonstrated that 27 wt% addition of Cat-8PEG in the PEG-oxime hydrogel system allows for greater retention on porcine cardiac tissue compared to Ald-AO. With this Ald-AO-Cat composition, the Ald-8PEG and Cat-8PEG are both available for binding to excess amines on the tissue surface for longer retention. Retention data also suggests that Cat-8PEG is retained in the gel via physical entrapment, which can help explain the longer retention on the *ex vivo* porcine tissue.

Prevention of cell adhesion is a major goal of this work, to ensure an efficient barrier is formed between the epicardium and other tissues to properly prevent adhesion formation within the first several weeks post-surgery.^4,57^ Following induced trauma, pericardial mesothelial cells detach from the native tissue layer as blood and inflammatory cells flow into the affected area. In the weeks that follow, the increased accumulation of fibrin, collagen and fibroblasts results in strong adhesive connections between the heart and surrounding tissue.^4^ PEG is well known to act as an anti-cell adhesion material and therefore we expected minimal cell adhesion on our oxime hydrogels.^58-60^ Cytocompatibility with L929 fibroblasts demonstrated the non-cytotoxic nature of Ald-AO-Cat. It was previously reported that Ald-AO oxime gels were cytocompatible, and therefore it was expected that similar results would be observed in this study. DA has been increasingly studied in recent years in biomaterials with adhesive properties, with similar results^39,43,61^ even though H_2_O_2_ is generated during catechol crosslinking.^62^ Therefore we did not expect Cat-8PEG to exhibit any cytotoxicity.

Typically, Cat-functionalized biomaterials are designed for tissue and cell adhesion applications.^39 42,52,63^ Conversely, we have successfully demonstrated the use of a catechol to prevent adhesions from forming between the epicardium and adjacent tissues in the chest cavity. We suspect that our incorporation of catechol functionalization (27 wt% Cat-8PEG) in Ald-AO-Cat ensures rapid binding of catechol to primary amines on the epicardial surface, and incorporation into the gel network, leaving limited unreacted catechol available for adhering to other tissue surfaces. To date, we have not seen a case of catechol used in this manner. From our previous unpublished *in vivo* studies, we determined that Ald-AO would not adequately be retained on the heart show efficacy as a cardiac antiadhesion barrier. By adding 27 wt% of catechol functionalization, we expected to see reduction of adhesion formation in the *in vivo* rat model. Re-entry into the chest after 2 weeks revealed reduced adhesion formation and intensity for Ald-AO-Cat compared to Ald-AO and untreated groups. The lower adhesion and intensity scores of Ald-AO-Cat at 2 weeks supported our claim that better retention and slower degradation allows for a more efficient cardiac anti-adhesion barrier system. Better efficacy for reducing adhesion formation and intensity was shown for rats treated with Ald-AO-Cat compared to Ald-AO, therefore only Ald-AO-Cat was studied at 4 weeks compared to untreated controls. At 4 weeks, complete reduction of adhesion formation was observed for Ald-AO-Cat rats, indicating its efficacy as a cardiac anti-adhesion barrier. Compared to the 2 week study, the overall lower adhesion scores reported at 4 weeks can be explained as a phenomenon commonly observed in the clinic. Generally, cardiothoracic surgeons report less pericardial adhesion coverage during re-sternotomy with increased waiting time prior to reoperation. As a limitation of the rat model, the rapid healing time of rats may have also contributed to low adhesion coverage observed at 4 weeks.

Echocardiography 3 days post-abrasion and material application did not show any signs of cardiac dysfunction. In previous reports of using CoSeal® in clinical trials, the 400 % swelling caused cardiac tamponade for several patients early in the study.^13,14^ Cardiac tamponade is a serious medical condition, resulting in compression on the heart and decreased cardiac function.^64^ Our oxime hydrogel was designed with 8-arm star PEG components, to allow for rapid gelation with many crosslinking sites, which ultimately results in minimal swelling and maintenance of systolic function.

Histological analysis was conducted at 4 weeks post material application to visualize any inflammation occurring on the epicardial surface in response to Ald-AO-Cat. Thin encapsulation occurred around the perimeter of the gel, which is expected for a foreign material implanted *in vivo.* As expected, we did not see any indication of cell infiltration into the gel, resulting from the anti-protein absorption properties of PEG.^58,59^ High resolution images of the capsule formation indicated some macrophage presence. This was also observed for CoSeal®, another PEG based polymer, in a porcine cardiac adhesion model.^1^ However, no lymphocytes or neutrophils were present with our oxime hydrogel, suggesting that Ald-AO-Cat did not cause chronic inflammation when administered on the epicardium. Given the thin encapsulation and lack of histological evidence of chronic inflammation this suggests that the Ald-AO-Cat does not impair the normal wound healing process. These findings are consistent with studies that showed biocompatibility of catechol functionalized materials used as tissue adhesives.^65,66^ Histological analysis also revealed that Ald-AO-Cat was not fully degraded up to 4 weeks post-delivery, providing a suitable barrier for reducing or eliminating the formation of cardiac adhesions. Cardiac adhesions are typically formed within the first 30 days after an open-heart procedure. The early and intermediate postoperative phases (1-30 days) are characterized by accumulation of fibrin, inflammatory cells and fibroblasts at the site of surgical trauma, and depressed fibrinolysis, allowing for the formation of fibrous tissue adhesions between the heart and adjacent tissues.^4,67^ Disrupting this process during the first 30 days post-op with a barrier device, is an effective way to prevent adhesion formation.

Finally, we developed a spray device that allowed simultaneous mixing and application of the two polymer solutions in a pilot porcine study. Arrangement of all channels for fluid flow in the assembled device ensured that airflow through each nozzle in the upper portion atomizes each liquid when pressure is applied to the syringes attached on the lower piece. Separation of the component solutions within the device is important due to the rapid gelation time of the PEG hydrogel which may impair delivery if mixing occurs prematurely. The overlapping trajectories of the droplets from both pairs of nozzles interact on the heart surface to form the gel layer. Testing of the spraying device indicated that adequate mixing of the two solutions occurred at a working distance of 12-17 cm leading to a homogenous gel formation and epicardial coverage, with minimal overspray, on the *ex vivo* heart. It is also important to note that with 4 mL of each solution being delivered, a 0.5-1 mm thick gel was formed, which mimics the application of CoSeal® in pre-clinical porcine models.^1^ Furthermore, the gel could be easily removed manually from the tissue surface, which would be important in the case a surgeon needs emergency re-access to the heart.

Compared to the non-treated controls, the pilot porcine studies suggest the use of Ald-AO-Cat as viable option for reducing the severity of cardiac adhesions in a clinical setting. The cardiac adhesion model that has been utilized here, consistently resulted in the formation of thick, fibrous adhesions that were difficult to remove and required sharp dissection. This is comparable to the adhesion model used previously to perform pre-clinical studies with CoSeal®, which resulted in nearly identical adhesion scores in the control group.^1^ Application of Ald-AO-Cat resulted in a reduction in adhesion severity, with the majority of the adhesions only requiring easy blunt dissection, and no tissue damage when they were removed. Compared to previous pre-clinical studies with CoSeal®, this pilot data suggests improved adhesion severity scores.^1^ It should be noted that in pediatric clinical trials, CoSeal® application was related to serious adverse events including cardiac tamponade, vena cava occlusion, and cardiac fibrillation, which were attributed to the large swelling capacity (400 % swelling) of CoSeal®.^14^ Here, we have mimicked the 8 mL dosing of CoSeal® used in the pre-clinical porcine study,^1,60^ and have shown indications of better reduction of adhesion severity, when scored on similar scoring systems. Because of the minimal swelling of Ald-AO-Cat compared to CoSeal® a future dosing study will allow us to determine if delivering larger amounts of oxime hydrogel will result in more favorable results, whereas clinical trials revealed that CoSeal® dosing was largely restricted by the inherent swelling behavior.^14^ As mentioned previously for the rat model, adhesion coverage also shows a decreasing trend from 3 weeks to 6 weeks in the porcine model, which is commonly observed in the clinic. Taken together, the rodent data along with the pilot pig data support the motivation for future dosing and efficacy studies of Ald-AO-Cat for cardiac adhesion prevention and reduction and continued translational development.

## Conclusion

In this study, we have provided evidence of a potential solution to the problem of post-surgical cardiac adhesions. When attempting to approach this condition using a physical barrier applied to the heart surface, it is important to consider the inherent properties of a proposed hydrogel in addition to its behavior when used in both *in vitro* and *in vivo* environments. We have demonstrated here that the Ald-AO-Cat hydrogel possesses mechanical characteristics and degradation kinetics that are well-suited to the conditions within the chest cavity. The material also demonstrated a lower degree of swelling that surpasses the behavior of existing products. The ability of this biocompatible Ald-AO-Cat hydrogel to prevent adhesion formation is demonstrated by reducing the onset of post-surgical adhesions *in vivo.* Pilot porcine cardiac anti-adhesion data supports further investigation of this hydrogel device for dosing and efficacy for potential future clinical translation.

## Acknowledgments

This work was funded by the National Heart, Lung, and Blood Institute UC CAI Grant U54HL119893 and National Center for Advancing Translational Sciences UCSD CTRI Grant 1UL1TR001442. G.M.P. was supported by an American Heart Association postdoctoral fellowship.

## Author Contributions

K.L.C. and M.M.M. obtained the funding. M.F., G.M.P., H.T.L., A.B., J.L.U., K.L.C., and M.M.M. designed the experiments. R.L.B. performed all small-animal surgeries and D.H. performed echocardiography. Data acquisition and/or analysis was performed by M.F., G.M.P., A.B., J.L.U., D.H., K.O., J.H.O., M.M.M., and K.L.C. M.F., G.L.P., and A.B. drafted the manuscript. Revising the manuscript for critical content was executed between by K.O., J.H.O., M.M.M., and K.L.C. Administrative, technical, or supervisory tasks were handled by K.L.C., R.L.B., and J.H.O.

## Methods

### Materials

Alcohol-terminated 8-arm PEG (molecular weight (MW 10 kDA) was purchased from Creative PEGWorks (North Carolina, USA). Alexa Fluor® 594 NHS Ester (Succinimidyl Ester) and Alexa Fluor® 594 Hydrazide were purchased from Thermo Fisher Scientific. All other chemicals were purchased from Sigma-Aldrich and were used as received without further purification.

### Synthesis of 8-arm aldehyde-PEG (Ald-8PEG) and 8-arm aminooxy-PEG (AO-8PEG)

Synthesis and purification of Ald-8PEG and AO-8PEG were performed as previously reported.^47.1^H NMR (Ald8PEG, 300 MHz, CDCl_3_) δ: 10.08 (s, 8H, C*H*O), 8.19 (m, 16H, *C_6_H_4_),* 7.93 (m, 16H, *C_6_H_4_),* 4.49 (m, 16H, C*H_2_*OCHO), 3.85-3.35 (m, C*H_2_*CH_2_OCHO and PEG protons). ^1^H NMR (AO8PEG, 300 MHz, D2O) δ: 3.88-3.85 (m, 2H, CH_2_ONH_2_), 3.8-3.4 (m, PEG protons). Ald-8PEG and AO-8PEG were stored under vacuum.

### Synthesis of 8-arm catechol-PEG (Cat-8PEG)

Synthesis of 4-nitrophenyl chloroformate (NPC)-activated 8-arm PEG (NPC-8PEG): Alcohol-terminated 8-arm PEG (MW 10 kDa) (5.00 g, 0.50 mmol) was dissolved in anhydrous dichloromethane (100 mL). The reaction flask was placed into an ice bath followed by addition of NPC (8.06 g, 40.00 mmol), trimethylamine (TEA) (5.58 mL, 40.00 mmol) and 4-(dimethylamino)pyridine (DMAP) (0.81 g, 4.00 mmol). After 3 h the reaction solution was poured into cold diethyl ether (900 mL), and the orange precipitate was collected via filtration and washed with cold diethyl ether. NPC-8PEG was obtained as a yellow solid. (5.65g, 99.81 % yield). ^1^H NMR (NPC8PEG, 300 MHz, CDCl3, δ): 8.26 (m, 16H, C6*H4*), 7.38 (m, 16H, C6*H4*), 4.43 (m, 16H, C*H_2_*OCOO), 3.85-3.35 (m, C*H_2_*CH_2_OCOO and PEG protons).

Synthesis of 8-arm catechol-PEG (Cat-8PEG): NPC-8PEG (5.65g, 0.50 mmol) was dissolved in anhydrous 1-methyl-2-pyrrolidone (113 mL), followed by addition of dopamine hydrochloride (7.57 g, 39.93 mmol) and TEA (5.57 g, 39.93 mmol). The reaction flask was kept at 60 °C for 12 h. The crude reaction product was dialyzed (MW cut off 3,500 g/mol) against water (pH = 3-4) to prevent the oxidative polymerization of DA, followed by filtration and lyophilization. Cat-8PEG was obtained as a solid (3.35 g) with 74 % functionalization according to ^1^H NMR. ^1^H NMR (Cat8PEG, 300 MHz, DMSO-d_6_) δ: 7.20 (m, 8H, N*H*), 6.59 (m, 8H, C6*H4*), 6.53 (m, 8H, C6*H4*), 6.39 (m, 8H, C6*H4*), 4.03 (m, 16H, C*H_2_*OCONH), 3.85-3.35 (m, C*H_2_*CH_2_OCONH and PEG protons). 3.40-3.05 (m, 32H, NHC*H_2_*CH_2_ and NHCH_2_C*H_2_*). Cat-functionalization was confirmed using a colorimetric DOPA assay developed by Waite and Benetict.^68^ In brief, a known amount of Cat-8PEG was dissolved in dilute acid. 100 μL of Cat-8PEG solution was added to 300 uL 0.5 M HCl in an Eppendorf tube. 300 μL of nitrite reagent (1.45 M sodium nitrite and 0.41 M sodium molybdate) and 400 μL of 1 M NaOH were added in rapid succession to the tube to form a red solution. Samples were read at 500 nm using a microplate reader (BioTek) within 2-5 minutes of NaOH addition. A standard curve was generated using known concentrations of 3,4-dihydroxyphenylalanine (*L*-DOPA) (Figure S4).

### Preparation of hydrogels

Hydrogels were prepared in water or 1X PBS, as specified for each experiment. In brief, for oxime hydrogels (Ald-AO), Ald-8PEG and AO-8PEG solutions were prepared and mixed to obtain final hydrogel solutions with a 1:1 ratio of the two polymer. For catechol-oxime hydrogels (Ald-AO-Cat), AO-8PEG solution was prepared separately from the Ald-8PEG and Cat-8PEG (Ald/Cat-8PEG) solution of the desired Cat-8PEG to Ald-8PEG ratio (Table 2). The two solutions were mixed to obtain a gel with the desired ratios of all three polymers. *In vivo,* each polymer solution was individually loaded into syringes and connected to a gas-assisted dual spray applicator tip (Nordson Medical, Loveland, Colorado).

### Characterization of hydrogels

Two polymer solutions (200 μL) (AO-8PEG and Ald-8PEG or Ald/Cat-8PEG) were added simultaneously to 4 mL glass vials and the timer started. As previously described, gelation time was measured at the inability of the solution to flow using a vial tilting method (n=3 for each formulation).^47,51^ Gelation was measured by hand for these gel formulations. Each hydrogel was incubated at 37 °C for 24 h and then placed into PBS pH 7.4 with 9-fold volume for 24 h. The swelling ratio (Q) was calculated as follows: *Q* = *V_s_/V_i_* * 100 (%), where *V_s_* is the volume of swollen hydrogels after 24 h and *V_i_* is the initial volume of hydrogels immediately after gelation. The swelling ratio of each hydrogel reached a plateau within 24 h and was defined as the equilibrium swelling ratio (n=3 for each formulation).

### Rheological studies

The storage modulus *(G’)* and loss modulus (*G*”) data were acquired using AR-G2 (TA Instruments) with a 20 mm parallel plate geometry operated at 37 °C by a frequency sweep from 10^−2^ to 10^2^ Hz (n=3 for each formulation). The hydrogels (500 μL, φ = 21 mm) were prepared on the stage of the rheometer and were measured with the 1200 mm gap between the plate and the stage of the rheometer after 15 minutes of hydrogel incubation. Relative G’ studies were conducted at 0, 1 and 3 hours post hydrogel gelation. Relative G’ was calculated as follows: Relative G’ = G’_t_ / G’_t=0_ * 100 (%), where G’_t_ is the G_t_’ at a given time and G’_t=0_ is the initial G’ after 15 min of gel incubation.

### *In vitro* degradation studies

The hydrogels (200 μL) were prepared in 4 mL glass vials and then placed into PBS pH 7.4 with 9-fold volume. PBS was replaced once a day and the hydrogels were weighed.

### *Ex vivo* hydrogel retention on cardiac tissues

As previously described.^47^ Ald-8PEG and AO8-PEG were respectively conjugated with Alexa Fluor® 594 dye. Biopsies of porcine heart within hours of euthanasia were taken with φ =5 mm punches. Each biopsy was dipped into PBS pH 7.4 to mimic the fluid present during a surgical procedure. The hydrogels were then formed using two pipettes (each 5 μL) resulting in 10 μL on the surface of the tissue. After 15 minutes, biopsies covered with hydrogels were placed into a 48 well plate with PBS (400 μL) containing 0.625 % vol/vol penstrep (Thermo Fisher Scientific) to completely cover the gel coated tissue. Buffer was immediately removed, and fluorescence was measured on a microplate reader (BioTek) (excite 589 nm, emission 615 nm, sensitivity 100), and fresh buffer was added (n=3 for each formulation). Cat-8PEG was not conjugated with the dye so fluorescence was derived from Ald-8PEG and AO-8PEG in the hydrogels.

### *In vitro* retention of Cat-8PEG in Ald-AO-Cat gels

Hydrogels were formed in 24-well plates with 100 μL Ald-Cat-8PEG in 1XPBS (200 mg/mL and 150 mg/mL) and 100 μL AO-8PEG in 1X PBS (200 mg/mL) (n=4). Gels were incubated at room temperature for 10 min. and were then washed with 1X PBS (500 μL) for 10 min on an orbital shaker at 50 rpm. The supernatant was collected (10 min time point), and new 1X PBS (500 μL) was added. Samples were incubated at 37 °C for up to 7 days, with 1XPBS changes each day. The supernatant was collected at 10 min, 1 d, 2 d, 3 d, and 7 d post-gelation to assess release of Cat-8PEG. Day 7 gels were collected at the end of the experiment. Cat-8PEG concentrated in the supernatant and day 7 gels was detected using a colorimetric DOPA assay as described previously.^68^ In brief, the supernatant samples were lyophilized and resuspended in 200 μL of dH_2_O. 50 μL of this solution was diluted 1:4 in 1XPBS. 100 μL 0.5 M HCl, 300 μL of nitrite reagent (1.45 M sodium nitrite and 0.41 M sodium molybdate) and 400 μL of 1 M NaOH were added in rapid succession to the Eppendorf tube to form a red solution. Samples were read at 500 nm using a Synergy 4 Multi-mode Microplate Reader (BioTek) within 2-5 minutes of NaOH addition. The same procedure was used for the day 7 gels (note: digestion of the gels does occur during this method, however complete digestion was not achieved, therefore trapping some Cat-8PEG in the remaining gel and reducing the absorbance reading.) A standard curve was generated using known concentrations of Cat-8PEG (Figure S5A).

### Inflammatory cell adhesion studies

The hydrogels (100 μL) were formed in a 96 well plate. The gels were swollen in Dulbecco’s PBS for 48 h at 37 °C. The membranes of L929 murine fibroblasts and RAW macrophages were labeled with PKH26 (Sigma, PKH26 Red Fluorescent Cell Linker Mini Kit for General Cell Membrane Labeling) following manufacture protocol. The PBS on the swollen gels was removed and either labeled L929 fibroblasts (100 μL, 200,000 cells/cm^2^) or RAW macrophages (100 μL, 100,000 cells/cm^2^) were seeded on top of the gels (n=5 for each formulation for each cell type); tissue culture plastic (TC) was used as a control. After 24 h at 37 °C, five fluorescent images were per well. The fluorescence per area was quantified using ImageJ.

## Direct Contact Assay

### Cell Culture

L929 murine fibroblasts were cultured with Dulbecco’s modified eagle’s medium (DMEM) supplemented with 10% fetal bovine serum (FBS) and 0.5% penicillin-streptomycin (P/S) and glutathione (2.61 mg/mL). Cells were allowed to grow to 70-90% confluency before plating. Cells were seeded 16,000 cells/cm^2^ and allowed to reach confluency (~24 h). Once confluent, baseline morphologic and metabolic activity were measured with bright field images and an alamar blue assay, respectively, for a baseline reading before treatment. The alamar blue assay was used as specified by the manufacturer’s protocol. After baseline measurements were obtained, three treatments were administered to the cells to test cytocompatabilty: 1) Ald-AO-Cat gel 2) positive control 3) negative control. Ald-AO-Cat gel solutions were prepared as described above and sterilized with a 0.22 μm filter before gelation. Gels were prepared directly on cell layers by combining 17 μL of each solution to the center of the cell layer and allowing 1 min for the material to gel completely and dry. Fresh culture media was added to the wells. Positive and negative controls were prepared by adding ZDEC (100mg/ml) and PBS (1:10, v/v) to the new media. After incubation for 24 hours at 37°C and 5% CO2, metabolic activity and cell morphology were measured as described previously. Micrographs of each sample set were taken for morphological analysis. In brief, micrographs taken from all groups were blinded prior to grading for healthy cell morphology, based on the criteria for biocompatibility outlined by the International Organization for Standardization (ISO 10993-5). These are outlined in the table below. Following grading, images were unblinded and the scores (**Table 3**) for each sample group were taken for statistical analysis.

### *In vivo* rat cardiac anti-adhesion ability of hydrogels

Cardiac anti-adhesion ability was evaluated on male Sprague-Dawley rats (Charles Rivers Laboratories Inc., San Diego, California) weighing 338 ± 28 g (3-4 months old). All study procedures were approved by the University of California, San Diego Institutional Animal Care and use Committee and performed in an American Association for Accreditation of Laboratory Animal Care (AAALAC) accredited facility. Animals were quarantined for at least 48 h after being received and were provided with food and water *ad libitum* before and after surgery. All animals were intubated and anesthetized using 5% isoflurane and subsequently shaved around the area of incision. Subjects were hydrated using a subcutaneous injection of lactated Ringer’s solution (3 mL) before lowering the isoflurane dosage to 2.5%. Animals were treated with 1% lidocaine administered subcutaneously.

Entry into the chest cavity was via thoracotomy. The pericardium was torn upon entry to access the surface of the heart. The tissue was then abraded by poking the exposed surface 100 times with a 30G syringe needle. This method was chosen for its consistent production in adhesion formation and severity after comparison to other techniques such as rubbing with a cytobrush, scalpel nicks and a decreased number of needle pokes. The heart was then allowed to dry via air exposure for 30 minutes before delivery of both hydrogel solutions (400 μL total) using a FibriJet gas-assisted applicator head from Nordson Micromedics. Test groups were treated to obtain 400 mL of either Ald-AO or Ald-AO-Cat gel over the tissue while an untreated group was used as a positive control. The gel was then allowed to dry for 10 minutes prior to closing of the chest cavity. Isoflurane dosage was lowered to 1% following closure of the ribs and turned off completely following final closure. Buprenex (0.05 mg/kg) was injected subcutaneously for pain relief. After 2 or 4 weeks, animals were euthanized via overdose Fatal Plus at 200 mg/kg. At 4 weeks, the hearts were dissected, paraffin embedded, sectioned, and stained with H&E. Samples from each group were blinded and graded by a histopathologist to assess the presence chronic inflammation.

### Dual-delivery device design and testing

A specialized device was developed to deliver each component together in a 1:1 ratio in the form of an atomized spray produced using compressed air and attached nozzle heads. This working prototype has two separate parts, one of which allows the flow of air through two separate pathways 0.4 cm in diameter after attachment to a compressor at an inlet while a second piece supports the addition of two 10 mL syringes containing sterile PBS solutions of the component polymers injected into the path of air through two pathways with a diameter of 0.2 cm. All nozzle heads have a diameter of 0.05 cm. The device was designed in SolidWorks and made with PLA filament on a Prusa i3 MK2 3D printer. The device was sterilized using EtO 48 hours before the procedure. For *in vivo use,* the device was operated at a gauge pressure of 10 psi at approximately 10-12 cm from the target surface while gel formation was visualized by adding India ink to each polymer solution (1:100 *v:v).*

The mixing of two solutions at a 1:1 ratio was confirmed with absorbance spectroscopy using dyed aqueous sugar solutions to mimic the viscosities of the AO-8PEG (0.01 Pa·s) and Ald-Cat-8PEG (0.04 Pa·s) solutions. Yellow dye with a peak absorbance at 420 nm was added to the PEG-AO substitute solution while a blue dye with a maximum at 610 nm was added to the PEG-Ald/PEG-DA substitute (2:25, v:v). A standard solution made from 250 μL of each dyed substitute was used as a control. The device was tested at working distances of 10 cm, 12.5 cm, 15 cm and 20 cm by spraying both solutions simultaneously into a petri dish 5.5 cm in diameter and collecting 500 μL of the final mixed solution. Absorbance spectroscopy performed using a TECAN plate reader measured the signal of each sprayed mixture at both 420 nm and 610 nm.

### *In vivo* porcine cardiac adhesion pilot study

Cardiac anti-adhesion ability was evaluated on male KG Farm Pig (S & S Farms, Ramona, California) weighing 15-27 kg (~1 month old) in a small pilot study. All study procedures were approved by the University of California, San Diego Institutional Animal Care and use Committee and performed in an American Association for Accreditation of Laboratory Animal Care (AAALAC) accredited facility. Animals were quarantined for at least 72 h after being received and were provided with food and water *ad libitum* before and after surgery. A Fentanyl patch (50 mcg) was placed for pain control 24 hours prior to median sternotomy. A combination of Ketamine (20-30 mg/kg)/Xylazine (2 mg/kg)/Atropine (0.04-0.05 mg/kg) was administered IM for pre-anesthesia induction. All animals were intubated and anesthetized with Propofol (2.4 mg/kg) with 1-3 % isofluorane administration maintained throughout the surgical procedure for deep anesthesia.

A median sternotomy (up to 20 cm incision) was performed to gain entry into the chest cavity. The pericardial sac was cut open and the heart was fully exposed. Adhesions induction was conducted by rubbing gauze 15-20 times on 5 regions of the anterior surface of the heart. 5 mL of blood was removed from the right atrium and sprayed on the epicardium and the chest cavity was left open to dry, exposed to air, for 15 minutes. Ald-Cat and AO solution in 1X PBS were prepared fresh and in a sterile hood prior to spraying. A total of 4 mL of each solution with India Ink (1:100) were prepared, filtered with a 0.22 μm sterile filter, and loaded into the spraying device. Fluid pooled in the chest cavity was aspirated prior to administering Ald-AO-Cat gel. Ald-AO-Cat gel (8 mL total) was sprayed onto the heart surface creating a 0.5-1 mm thick gel (working pressure of 10 psi and a working distance of ~12-17 cm). After spraying, the chest was left open, exposed to air for 10 minutes before closing. Control animals received the same treatment, without the spray of a gel. The pericardium was left open, sternal wires were used to close the chest, and sutures were used to for closing subcutaneous and skin layers. Fentanyl (10 mcg/kgm, IV), Baytril (10mg/kg, IM) and Flunixin (2 mg/kg, IM) were administered intra-operatively and Bupivicain was injected at the incision site before wrapping the chest with gauze. Fentanyl (5 mg/kg) was delivered IV during recovery. Carprofen (2 mg/kg, IM, SID) for 3 days post-op and Clavamox (375 mg, PO BID) was delivered for 5-7 days post-op.

Pigs were euthanized at 3 and 6 weeks post sternotomy with an overdose of Fatal-Plus (9 mL). Pigs that received Ald-AO-Cat spray were euthanized at 3 (n=1) and 6 (n=1) weeks post sternotomy. The control pigs (n=3) were euthanized 6 weeks post sternotomy. Each animal was euthanized followed by resternotomy and adhesion gross assessment. Images of the anterior hearts surface immediately following resternotomy were used for blinded adhesion grading of coverage. Adhesion strength was assessed during resternotomy. Images from resternotomy were segmented into 6 equivalent sections (similar to the rat grading criteria), and each section was graded on a scale of 1-4 for adhesion coverage and strength as shown in Table 4. The scoring system was adapted from several surgical adhesion studies.^48’50,60,69,70^

## Statistical Analysis

Statistical differences between the experimental data for the 4 week *in vivo* adhesion assessment were analyzed using a Mann-Whitney t-test. All rheological data, cell adhesion and cytocompatibility, 2 week *in vivo* data were analyzed using one-way analysis of variance (ANOVA) with Tukey’s post hoc analysis. Statistical significance was set to p < 0.05. Results are presented as means ± standard deviation.

## Availability of Data

The data that support the findings of this study are available from the corresponding author upon reasonable request.

