## Supplemental Info for "Preventing post-surgical cardiac adhesions with a catechol-functionalized oxime hydrogel"

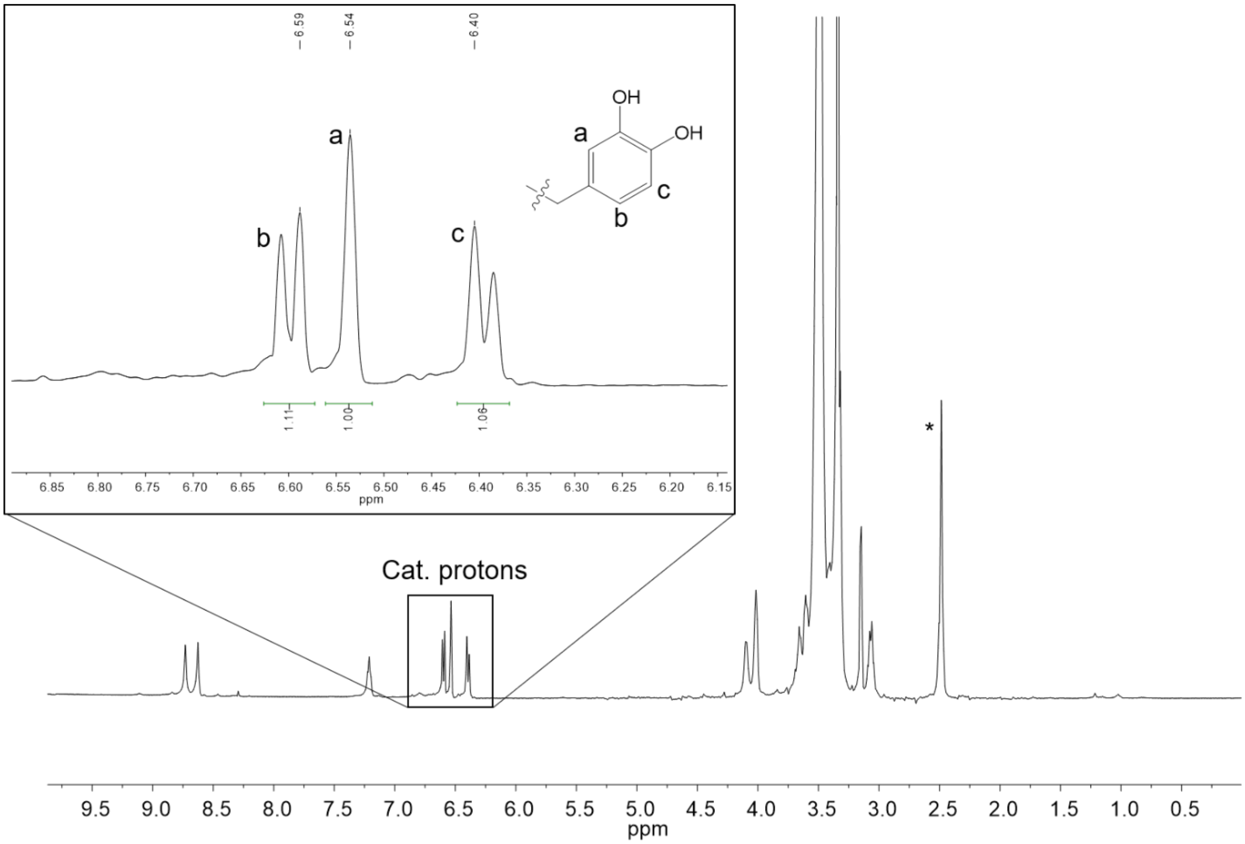


**Figure S1.** $^1$H NMR was used to determine the successful synthesis of Cat-8PEG and determine the functionality ratio. Characteristic peaks at δ= 6.59 ppm, 6.54 ppm and 6.40 ppm are representative of the DOPA catechol group functionalization. Cat8PEG was obtained as a solid (3.35 g) with 74 % functionalization. $^1$H NMR (Cat8PEG, 300 MHz, DMSO-$d_6$ (*)) δ: 8.73 (s, 8H, O*H*), 8.63 (s, 8H, O*H*), 7.20 (m, 8H, N*H*), 6.59 (m, 8H, $C_6H_4$), 6.53 (m, 8H, $C_6H_4$), 6.39 (m, 8H, $C_6H_4$), 4.10-4.02 (m, 32H, $CH_2CH_2$OCONH), 3.85-3.35 (m, PEG protons). 3.15-3.05 (m, 32H, NH$CH_2CH_2$). Cat-8PEG was stored under vacuum.

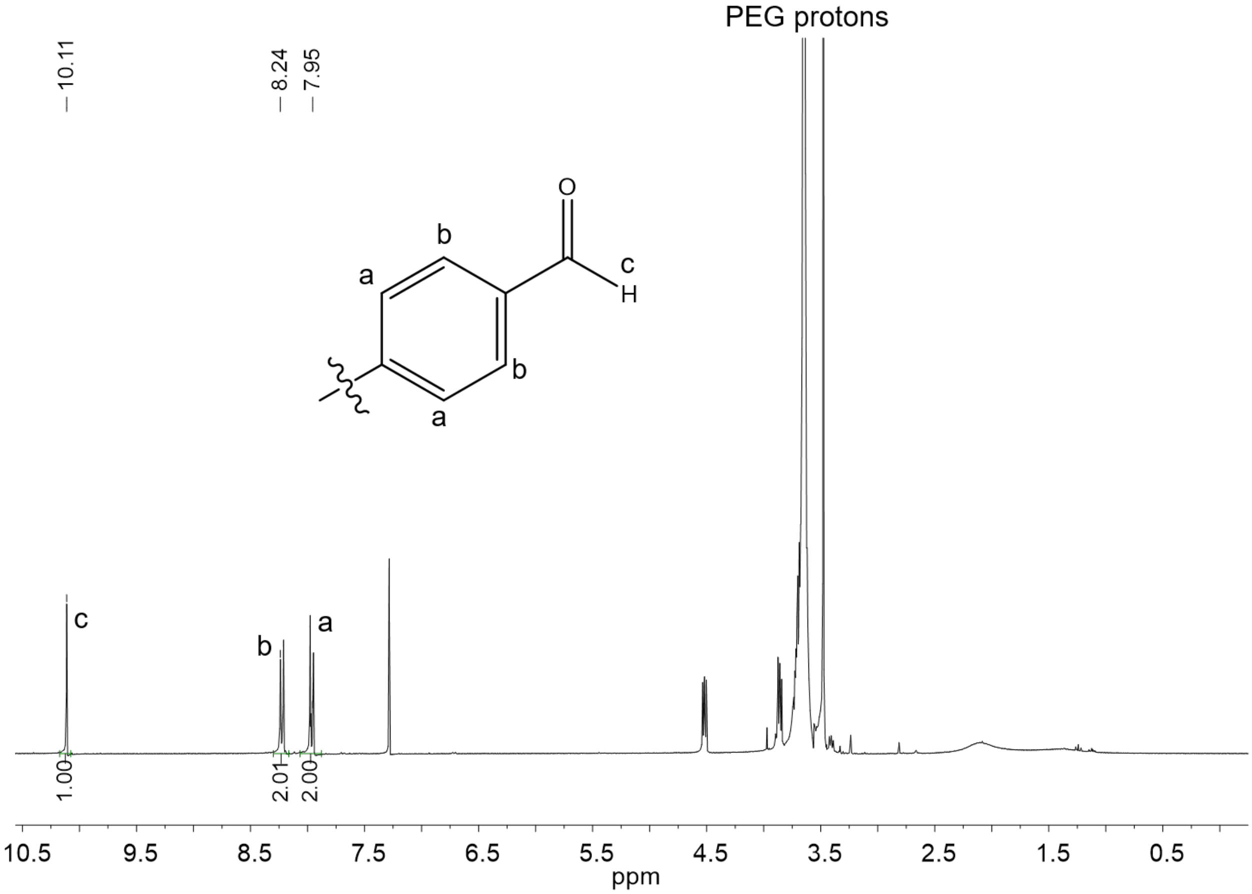


**Figure S2.** $^1$H NMR was used to determine the successful synthesis of Ald-8PEG and determine the functionality ratio. Characteristic peaks at δ= 10.11 ppm, 8.24 ppm and 7.95 ppm are representative of the benzaldehyde functionality. Ald-8PEG was obtained as a white solid with 97 % functionalization. $^1$H NMR (Ald-8PEG, 300 MHz, $CDCl_3$) δ: 10.11 (s, 8H, C*H*O), 8.24 (m, 16H, $C_6H_4$), 7.95 (m, 16H, $C_6H_4$), 4.49 (m, 16H, $CH_2OCHO$), 3.85-3.35 (m, $CH_2CH_2OCHO$ and PEG protons). Ald-8PEG was stored under vacuum until use.

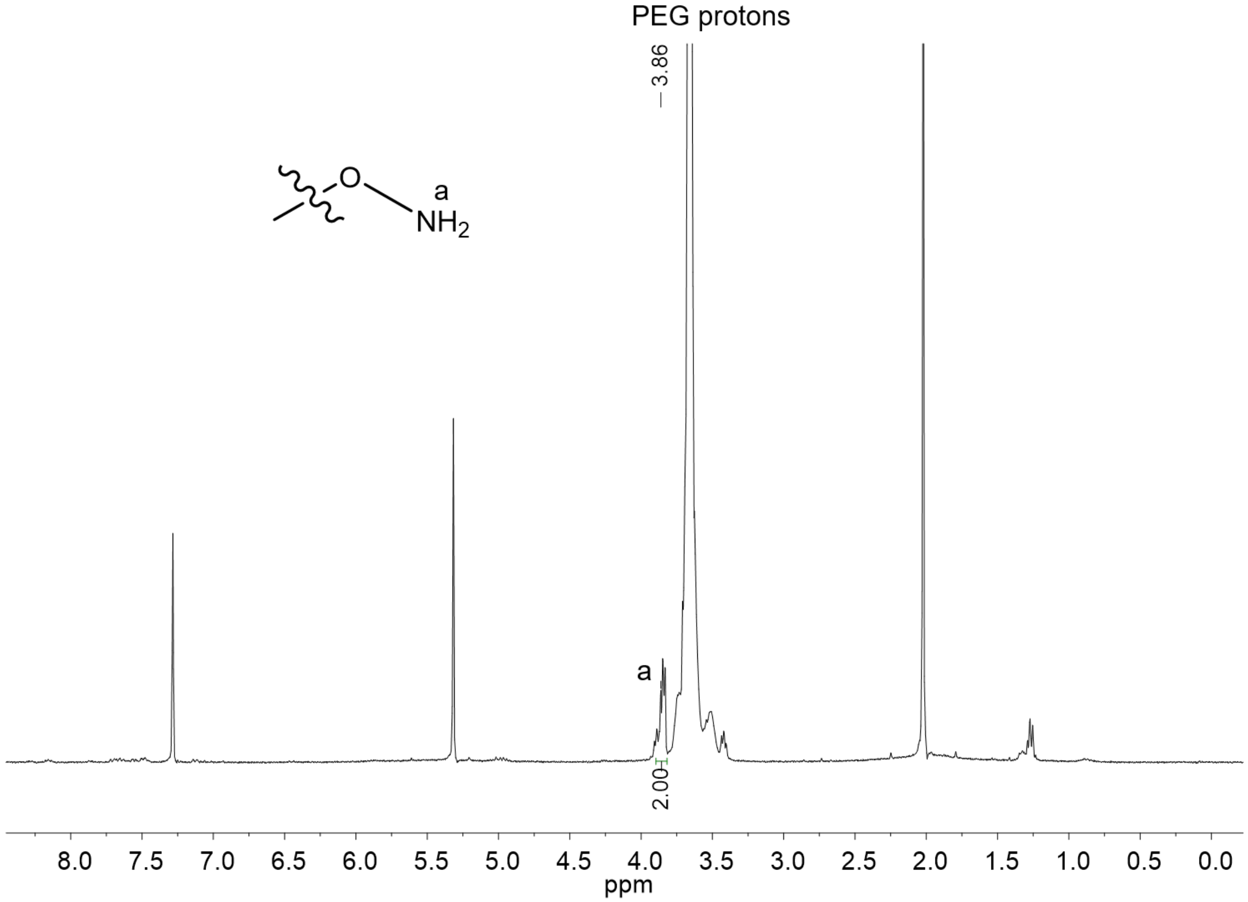


**Figure S3.** [1]H NMR was used to determine the successful synthesis of AO-8PEG and determine the functionality ratio. The characteristic peak at δ= 3.86 ppm represents the aminooxy functionality. AO-8PEG was obtained as a white solid with 83 % functionalization. [1]H NMR (AO-8PEG, 300 MHz, $D_2O$) δ: 3.86 (m, 2H, CH2ONH2), 3.8-3.4 (m, PEG protons). AO-8PEG was stored under vacuum until use.

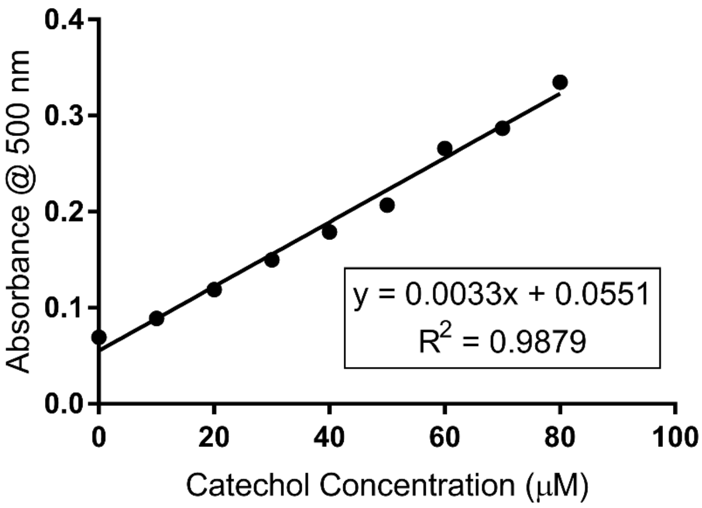


**Figure S4.** A standard curve was generated using known concentrations of 3,4-dihydroxy-L-phenylalanine for quantifying catechol functionalization in Cat-8PEG. Absorbance was recorded at 500 nm.

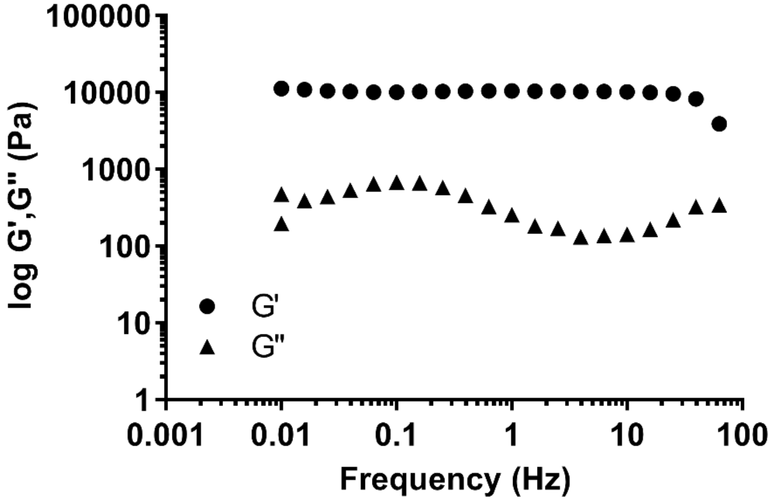


**Figure S5.** Representative G' and G" across a frequency sweep. This graph is representative of the viscoelastic behavior of both the Ald-AO and Ald-AO-Cat gels which crosslinked and gelled within seconds of combining the two solutions prior to rheometry.

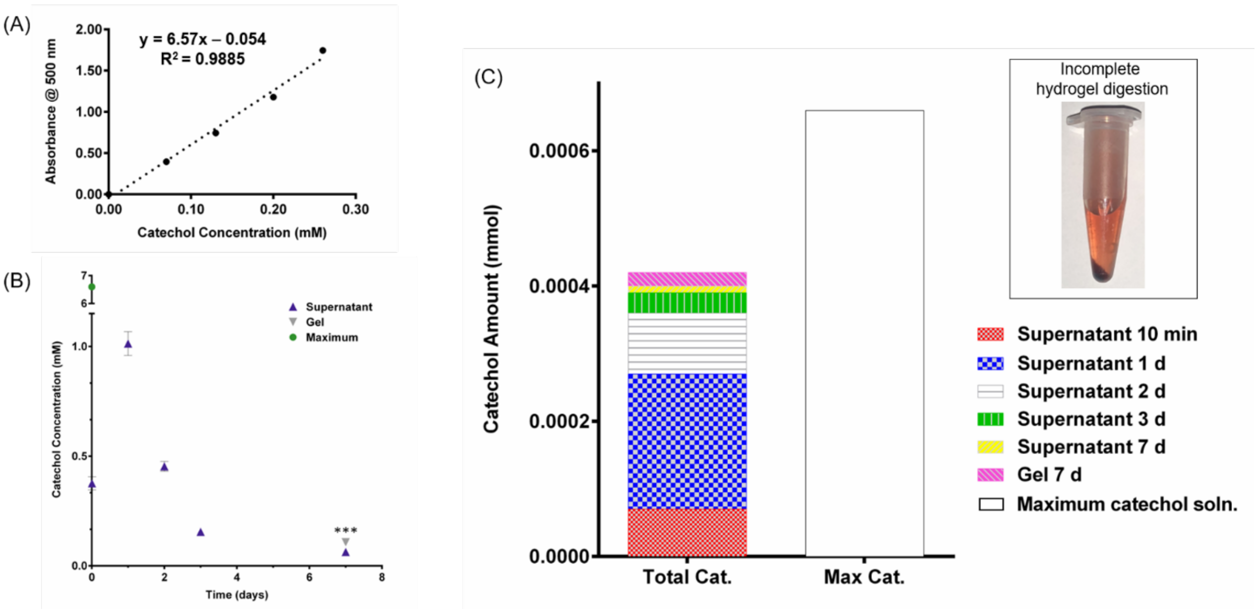


**Figure S6.** Catechol is retained in the Ald-AO-Cat as indicated by the time-dependent DOPA-nitration assay. (A) Release of Cat-8PEG was detected in gel supernatant at 10 min, 1 day, 2 days, 3 days, and 7 days post-gelation. An initial release of Cat-8PEG was observed at 10 min, corresponding to the release of Ald-8PEG and AO-8PEG observed in the ex vivo retention assay (Figure 2F). This is indicative of the release of unreacted or non-physically trapped materials. A burst release of Cat-8PEG was detected, similar to the release of Ald-8PEG and AO-8PEG, which was expected with maximum swelling of the oxime hydrogel in aqueous conditions. A time-dependent, decrease in Cat-8PEG release was calculated up to day 7, with a significantly larger amount of Cat-8PEG physically trapped in the gel at 7 days post gelation. (B) The summation of Cat-8PEG released in the supernatant and retained in the gel at day 7, was compared to the total Cat-8PEG concentration in the original gel formulation, with 63 % of the total Cat-8PEG accounted for. The remaining Cat-8PEG can be explained by the incomplete digestion of the gels at day 7 resulting in incomplete release of Cat-8PEG for detection.

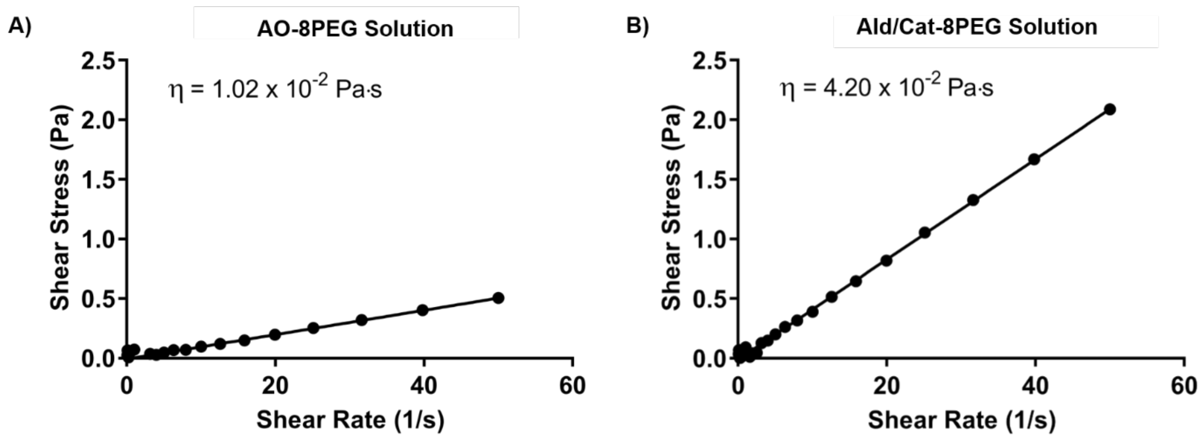


**Figure S7.** Parallel plate rheometry was used to measure the viscosity (η) of the Ald-8PEG/Cat-8PEG (A) and AO-PEG (B) solutions.

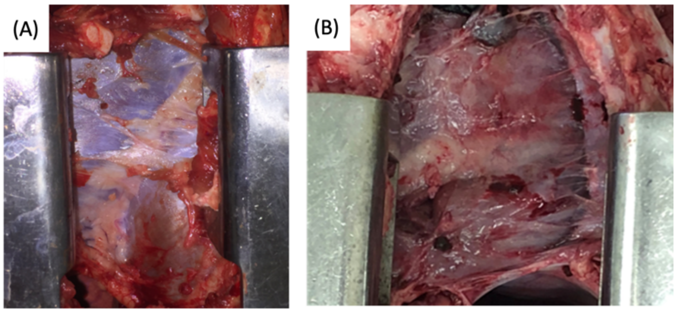


**Figure S8.** Representative images from pilot pig study showing robust adhesions in the control group (A) that required sharp dissection, whereas Ald-AO-Cat treated pigs (B) had milder adhesions that required only manual/blunt dissection. B displays Ald-AO-Cat treated pig euthanized at 3 weeks after application.
